# A Unified World Model in Human Posterior Parietal Cortex

**DOI:** 10.64898/2026.10.30.762936

**Authors:** V. Bougou, R. T. Pramod, E. R. Rosario, K. Pejsa, A. Bari, J. B. Tenenbaum, N. Kanwisher, R. A. Andersen

## Abstract

How does the brain understand the physical world, predict future events, and plan actions? Are these problems solved by a unified “world model,” as hypothesized by cognitive scientists and artificial intelligence researchers? Posterior parietal cortex has been implicated in all these functions, but whether the same neurons support each of them is unknown. We recorded from the superior parietal lobule (SPL) of two tetraplegic participants implanted with microelectrode arrays. SPL units preferentially engaged by intuitive physics—judging block-tower stability and viewing naturalistic physical events—also predicted whether moving objects would collide hundreds of milliseconds before contact, generalizing between different physical scenarios. These populations also encoded which hand action the participant intended to perform. These findings identify SPL as a substrate for physical world modeling.

## Introduction

Adaptive behavior requires continuously interpreting a changing physical environment, predicting how objects move and events unfold, and planning our own actions in response, often all within a fraction of a second. Cognitive scientists and theorists of motor control have long proposed that this capacity relies on internal models of the world, and building such world models in machines is a central aspiration of current research in artificial intelligence (*1–5*). Recent frameworks describe three core functions of a unified world model: i) building a physical model of the scene, including representations of discrete objects, their physical properties, and the forces acting upon them, ii) running simulations to predict future states, and iii) planning action (*5–8*) (Fig. 1A). Here we ask whether these three functions draw on a common neural substrate in the human brain.

**Fig. 1.**
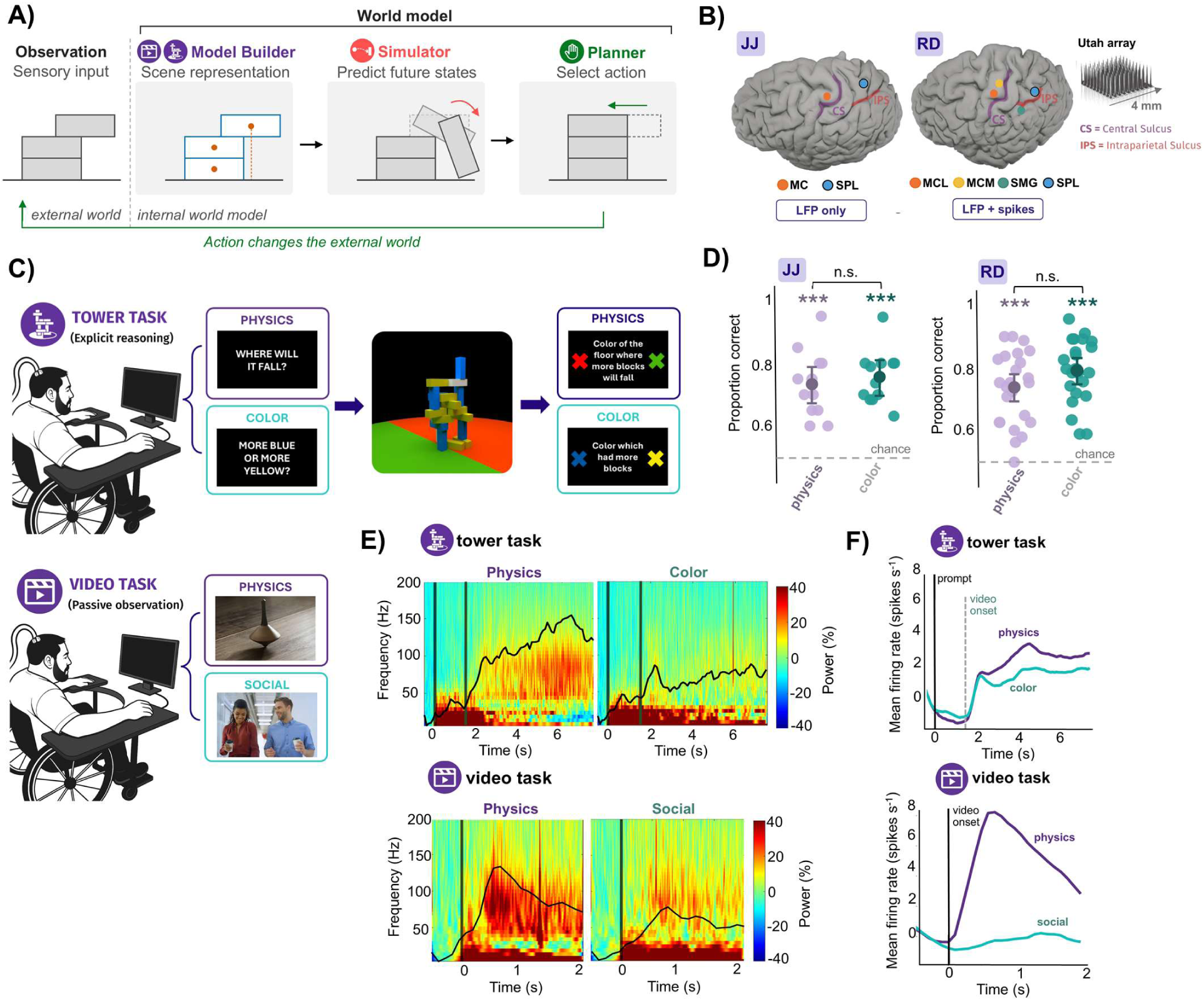
Experimental design and neural responses across physical reasoning tasks. **(A)** World model framework with three functions: i) a model builder that uses sensory input from the external world to construct an internal physical model of the scene including discrete representations of individual objects, their physical properties, and the forces acting upon them, ii) a simulator that predicts future states (successive frames of the falling block), and iii) a planner that selects an action (re-centering the block). Actions change the world, closing the loop. Icons at top indicate the four experiments testing each function. **(B)** Intracortical recording locations for participants JJ and RD. **(C)** Experimental paradigms. Participants completed a tower task requiring either explicit physical reasoning about tower stability (physics condition) or a perceptual judgment about block color (color control), using identical visual stimuli. Participants also passively viewed naturalistic videos depicting either physical interactions (physics condition) or social interactions (social control). **(D)** Behavioral performance during the tower task for JJ (left) and RD (right). Pale jittered points show accuracy for individual sessions for physics and color judgments. Large circles indicate accuracy pooled across all trials of each condition, with error bars showing 95% Clopper–Pearson exact binomial confidence intervals. Performance was significantly above chance for both conditions in both participants (***P < 0.001), with no significant difference in accuracy between physics and color judgments (n.s.). **(E)** Time-frequency representations of LFP power in JJ during the tower task (top) and video task (bottom), averaged across trials and recording channels. Physics conditions are shown on the left and their respective color or social conditions on the right. Color indicates change in spectral power relative to baseline. Black traces show high-gamma (60–120 Hz) power. **(F)** Population-averaged firing rates in RD during the tower task (left) and video task (right), averaged across trials and units significantly modulated in at least one condition. Vertical dashed lines indicate relevant task events.

The posterior parietal cortex (PPC) is a compelling candidate for such a substrate. Classic lesion and neuroimaging studies have established a central role for parietal cortex in visually guided action (*9*, *10*), while neurophysiological studies in nonhuman primates and humans have demonstrated representations of action goals and intentions in PPC (*11–13*). Separately, human neuroimaging studies (*14–16*) have identified a set of parietal regions that are selectively engaged during intuitive physical reasoning, termed the “physics network”. These regions respond when people reason about the stability, mass, and interactions of objects, suggesting that PPC represents information about the physical structure and dynamics of the environment. Although these findings implicate nearby and potentially overlapping cortical areas in understanding the physical world and planning actions within it, it remains unknown whether these functions are supported by the same neuronal population as an integrated world model would predict.

Here, we use intracortical recordings from microelectrode arrays implanted in two individuals with tetraplegia to test the hypothesis that the three core functions of a physical world model engage the same neural population (Fig. 1A) in human superior parietal lobule (SPL). Two experiments isolate the building of an internal physical model of the scene, a third experiment tests whether the same neural population predicts physical events before they occur, and a fourth experiment tests whether the same neural population holds action intentions. We find that the same neural population, and even some of the same individual units, exhibit all three signatures of a physical world model. These findings provide neuronal-level evidence that physical reasoning, prediction, and action planning draw on a common representational substrate in human SPL, thus identifying this region as a cortical locus of the long-hypothesized world model in the human brain (*1*).

## Results

### Participants and recordings

We recorded single- and multi-unit activity as well as local field potentials (LFPs) from two tetraplegic participants (JJ, RD) implanted with Utah arrays targeting motor and posterior parietal cortices. Participant JJ had 96-channel arrays in the hand knob area of primary motor cortex and SPL. Participant RD had 64-channel arrays in medial and lateral motor cortex, SPL, and supramarginal gyrus. Here, we focus on activity recorded from the SPL arrays in both participants, as fMRI studies have identified it as a node in the physical reasoning network.

Unit counts for each experiment are reported in tables S1 to S3; implant locations are shown in Fig. 1B. Across experiments, we recorded 1,102 sorted units from SPL in participant RD across 12 sessions (unit-sessions; see fig. S1 for re-identification), of which 234 were well isolated (quality 1–2, see Materials and Methods), with the remainder classified as multiunit activity (quality 3–4). In JJ, whose arrays had been implanted for an extended period, we report only LFP activity.

Because recordings were obtained from the same arrays across multiple sessions, we evaluated potential re-identification of units across days in RD. The median percentage of same-electrode unit pairs classified as putative cross-session matches was 7.8% (range, 0.0–38.9%; fig. S1), suggesting limited cross-session overlap on average.

### Neural responses show consistent selectivity for intuitive physical reasoning across tasks

We first asked whether intracortical recordings from SPL showed the preferential responses to physics tasks previously shown with fMRI (*14*). Because the anatomical location of the fMRI-defined physics network varies across individuals and each Utah array samples only a small cortical region (4 × 4 mm), we tested for physics-selective responses at the implanted SPL sites using the tower task (*14*), previously used to localize the physics network in fMRI studies (Fig. 1C). We then examined whether physics selectivity extended to passive viewing of naturalistic videos depicting physical or social interactions (“video task”; Fig. 1C).

In the tower task, participants judged whether a rotating tower of blocks would fall toward the red or green side of the floor (physics condition) or contained more blue or yellow blocks (color condition), a difficulty-matched perceptual judgment using the same stimuli. Both participants performed significantly above chance, with no significant difference between conditions (Fig. 1D). Eye tracking showed comparable gaze distributions across conditions and tasks (fig. S2A).

Both participants showed preferential LFP responses to physics conditions in both tasks. In JJ, high-gamma power was significantly greater during physics than color trials in the tower task, emerging 2.5 s after video onset, corresponding to 4 s after prompt onset (23.7% vs. 9.6% change from baseline during the significant epoch; unpaired two-sample t-test, Bonferroni-corrected across windows; Fig. 1E). Similarly, in the video task, high-gamma power was greater during physical than social videos, emerging 325 ms after onset (23.9% vs. 11.1%; unpaired two-sample t-test, Bonferroni-corrected). RD showed similar LFP responses in both tasks (fig. S2, B and C).

Neuronal firing rates in RD showed a corresponding preference for physics (Fig. 1F). In the tower task, population activity was significantly greater during physics than color trials (2.67 vs. 1.53 spikes/s during the significant epoch, emerging 1.4 s after video onset, corresponding to 2.9 s after prompt onset; unpaired t-test, Bonferroni-corrected across time windows). This preference was more pronounced in the video task, with increased firing during physical videos but near-baseline activity during social videos (5.12 vs. −0.26 spikes/s, emerging 100 ms after video onset). Median response latencies were 2.3 s after prompt onset in the tower task (800 ms after video onset) and 500 ms after video onset in the video task (Fig. 2A).

**Fig. 2.**
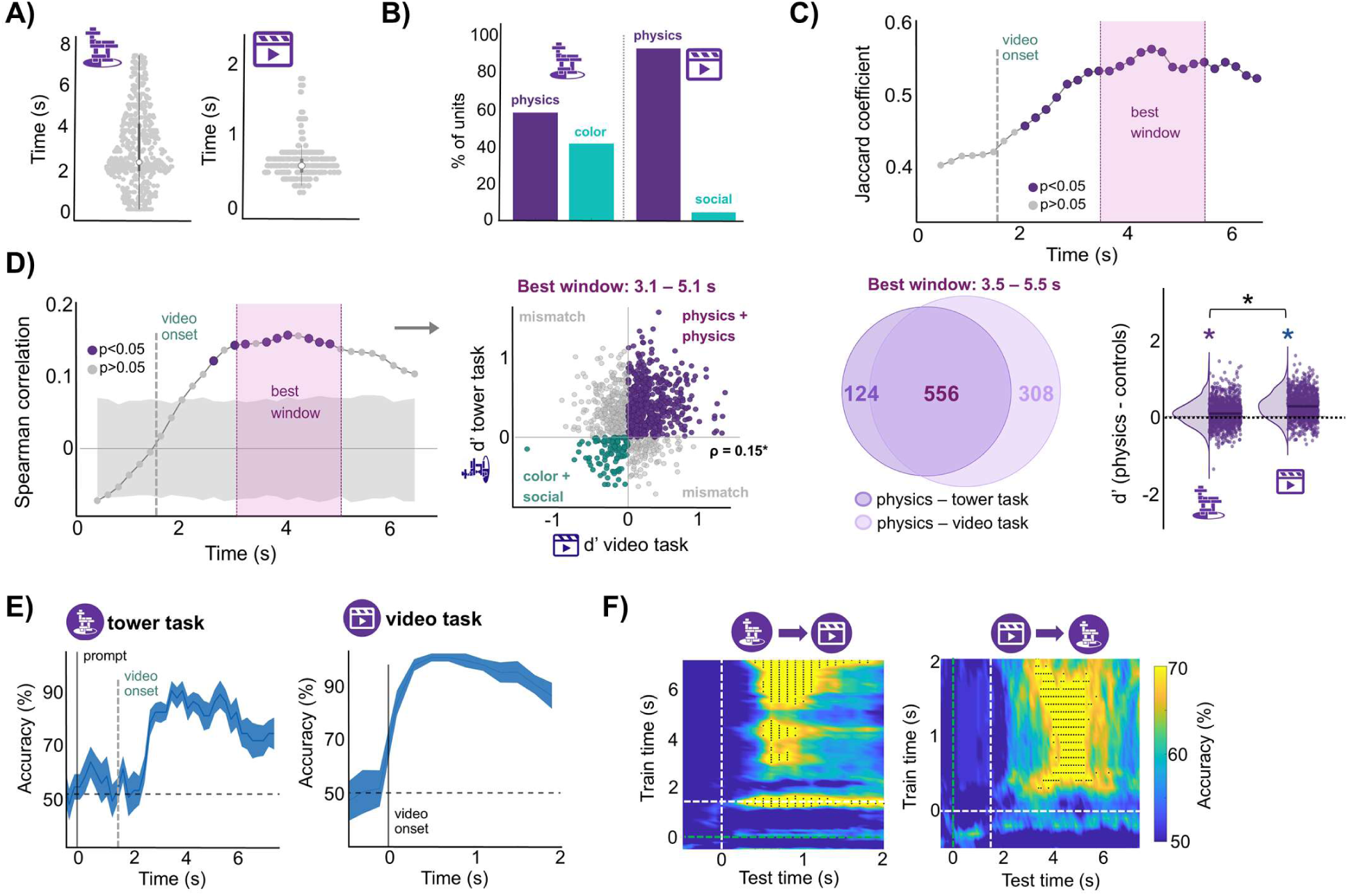
Shared neural representations of physics across tasks. **(A)** Response latencies of individual units in RD during the tower task (left) and video task (right) relative to prompt onset and video onset respectively. **(B)** Percentage of units in RD showing a significant response relative to baseline during the physics and control conditions of the tower task (left) and video task (right). **(C)** Overlap of physics-preferring units (d′ > 0) between the tower and video tasks. Top: Jaccard index as a function of the tower-task analysis window. Purple points indicate significant overlap relative to a label-permutation null (Bonferroni-corrected *P* < 0.05); gray points indicate nonsignificant overlap. The dashed vertical line marks tower-video onset. Bottom left: overlap at the window with the highest significant Jaccard index (3.5–5.5 s after prompt onset). Bottom right: distributions of unit-level d′ values during this window. The dashed line marks d′ = 0. Asterisks indicate significant differences from zero or between tasks (two-sided Wilcoxon signed-rank tests, *P* < 0.05). **(D)** Left: Spearman correlation between unit-level d′ values in successive 2-s tower-task windows and the full 2-s video-task period. Purple and gray points indicate significant and nonsignificant correlations, respectively (label-permutation tests, Bonferroni-corrected *P* < 0.05). The gray band shows the pointwise 5th–95th percentiles of the permutation distribution. The shaded region marks the window with the highest significant correlation (3.1–5.1 s after prompt onset). Right: Correspondence between video-task and tower-task d′ values during the best window. Each point represents one unit. Purple indicates physics preference in both tasks, teal indicates control preference in both tasks, and gray indicates opposite preferences across tasks. ρ = Spearman correlation for the best window (ρ = 0.15, *P* < 0.05). **(E)** Cross-validated LDA decoding of physics versus control conditions within the tower task (left) and video task (right). Lines and shading show mean accuracy ± SEM across cross-validation folds. Tower-task time is relative to prompt onset; solid and dashed vertical lines mark prompt and video onset, respectively. Video-task time is relative to video onset. Horizontal dashed lines indicate chance performance (50%). **(F)** Cross-task generalization of physics decoding. Classifiers trained at each time window in the tower task were tested across time in the video task (left), and classifiers trained in the video task were tested across time in the tower task (right). Color indicates physics-versus-control decoding accuracy. Black dots indicate train–test time pairs with significant above-chance decoding based on permutation testing with maximum-statistic correction across the full matrix (family-wise corrected *P* < 0.05). Dashed lines mark task events (green: prompt onset, white: video onset).

The prevalence of task-responsive units showed a similar pattern (Fig. 2B), with 58.3% responding significantly to physics versus 41.6% to color trials in the tower task, and 93.7% to physical versus 6.2% to social videos in the video task. Responses were predominantly excitatory in both tower conditions (60.6% for physics and 54.6% for color), whereas 81.9% of units showed excitation during physical videos and 53.2% showed suppression during social videos (fig. S2D).

To test whether the presence of visible body parts or intentional actions could account for physics selectivity in the video task, we compared responses to physical videos with and without each feature (fig. S2, E and F; see Materials and Methods). Neither population responses nor selectivity indices showed systematic differences between subcategories, providing no evidence that either feature explained the preference for physical videos.

Together, these results provide evidence for a physics-selective population in human SPL, extending previous fMRI findings by demonstrating consistent physics-related responses at the level of neuronal populations across explicit physical reasoning and passive observation of physical events.

### Physics-selective representations generalize across tasks

We next asked whether physics preference across tasks emerged from the same neuronal populations, a question that cannot be resolved with fMRI. For each unit, we calculated a selectivity index (d′) contrasting physics and control conditions, with positive values indicating physics preference. We quantified the overlap of physics-preferring units (d′ > 0) across tasks using the Jaccard coefficient (Fig. 2C). Overlap was significant relative to chance in most windows, peaking at 0.56 between 3.5 and 5.5 s from prompt onset (2–4 s after tower video onset). At this peak, d′ distributions were significantly shifted above zero in both tasks (tower: median d′ = 0.08; video: median d′ = 0.27; both *P* < 0.05, two-sided Wilcoxon signed-rank tests), with stronger selectivity in the video task (*P* < 0.05, paired Wilcoxon signed-rank test). During this window, 556 units preferred physics in both tasks, compared with 124 tower-only and 308 video-only units, representing 81.8% and 64.4% of physics-preferring units in the respective tasks. Example neurons are shown in fig. S3.

Beyond this categorical overlap, we examined whether the magnitude of unit-level selectivity was preserved across tasks. Selectivity indices were significantly correlated, peaking between 3.1 and 5.1 s in the tower task (Spearman’s ρ = 0.15, *P* < 0.05; Fig. 2D). Correcting for within-task measurement reliability increased the estimated correlation to ρ = 0.30 (IQR [0.29, 0.32]; see Materials and Methods), suggesting that measurement variability partly limited the observed correlation. Thus, units with stronger physics preferences in one task tended to show stronger preferences in the other.

At the population level, neural activity reliably discriminated physics from control conditions, reaching peak decoding accuracies of 86.25% in the tower task and 98.6% in the video task (Fig. 2E). To determine whether the high decoding accuracy in the video task was driven by suppressive responses to social videos, we repeated the analysis after excluding social-suppressed units. Decoding remained high (94.6% vs. 98.6% with all units; fig. S4A). Cross-temporal decoding revealed stable representations across extended periods of both tasks (fig. S4B). Decoding also remained above chance across individual sessions and was largely unchanged when excluding each session in turn, indicating that the effects were consistent across recording days (fig. S4C). Accuracy showed no systematic increase or decrease across sessions, arguing against learning across recording days or a transient attentional effect driven by novelty.

Critically, decoding generalized across tasks despite the different stimuli, behavioral demands, and control conditions (Fig. 2F). A decoder trained to distinguish physics from color trials in the tower task successfully discriminated physical from social videos, reaching a maximum significant accuracy of 91.7%. Generalization was also observed in the opposite direction, with a decoder trained on the video task reaching 77.5% accuracy when tested on the tower task. Thus, the population-level distinction between physics and control conditions was sufficiently similar across tasks to support bidirectional cross-task decoding.

Finally, repeating all analyses using only well-isolated single units yielded the same pattern of results (fig. S5), confirming that the observed effects did not depend on the inclusion of multiunit activity.

Together, these results show that physics selectivity generalizes across tasks at both the single-unit and population levels. Many of the same units preferentially responded during explicit judgments of physical stability and passive observation of physical events (see fig. S3 for examples), while the population response patterns evoked by the physics conditions were aligned across tasks. This convergence is particularly notable given that the two tasks differed in their visual input, behavioral demands, and control conditions, implicating the SPL in the construction of a physical model of the scene.

### Tuning of individual units reveals encoding of predicted outcomes

A second central function of an internal world model is to predict future physical states. We therefore asked whether SPL activity encoded the outcome of upcoming physical events. Participant RD passively viewed videos in which a moving object either collided with another object (hit) or missed, across two visually distinct scenarios in which the object was either rolled or thrown (Fig. 3A). Neural responses were aligned to contact time, which varied across videos (see Materials and Methods). Gaze distributions are shown in fig. S6A.

**Fig. 3.**
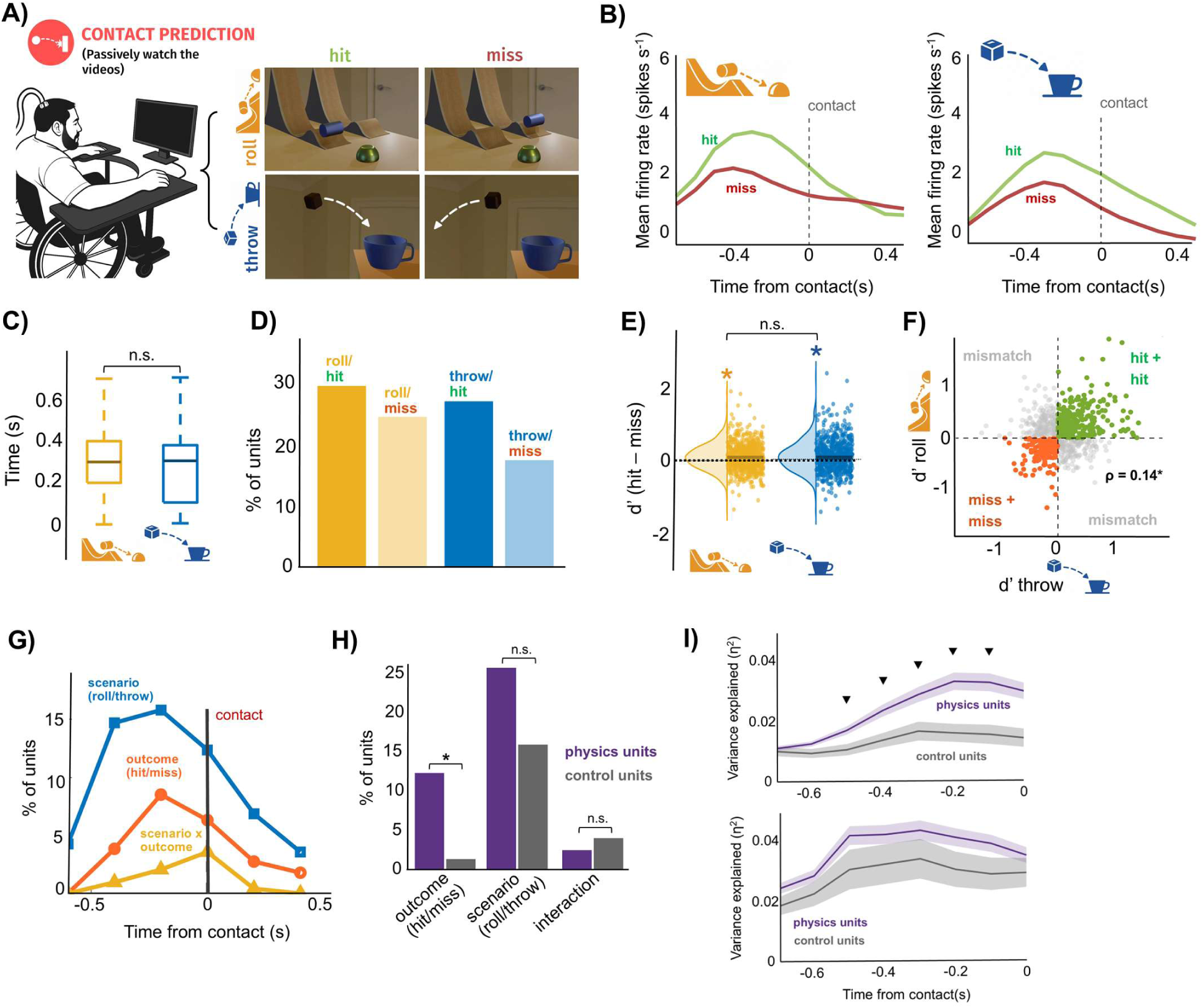
Neural encoding of physical outcomes across scenarios. **(A)** Contact-prediction task. Participant RD passively viewed videos depicting two physical scenarios, roll and throw, in which an approaching object either contacted the target (hit) or missed it (miss). **(B)** Population-averaged firing rates aligned to contact (t = 0) for the roll (left) and throw (right) scenarios, shown separately for hit (green) and miss (red) trials. **(C)** Distribution of response latencies (time of peak firing relative to contact) across units for roll (orange) and throw (blue) scenarios. n.s. indicates no significant difference between scenarios. **(D)** Percentage of units showing a significant response relative to baseline for each scenario and outcome: roll/hit, roll/miss, throw/hit, and throw/miss. **(E)** Outcome selectivity across scenarios. Distributions of unit-level d′ for hit versus miss trials are shown for roll (orange) and throw (blue). Asterisks indicate distributions significantly different from zero (*P* < 0.05); n.s. indicates no significant difference between scenarios. **(F)** Correspondence of unit-level outcome selectivity across scenarios. Each point represents one unit, with outcome d′ during throw trials on the x-axis and roll trials on the y-axis. Green indicates units preferring hits in both scenarios, red indicates units preferring misses in both scenarios, and gray indicates units with opposite selectivity across scenarios. Outcome selectivity was positively correlated across scenarios (Spearman ρ = 0.14, *P* < 0.05). **(G)** Time-resolved single-unit selectivity for scenario and outcome. Percentage of units showing a significant main effect of scenario (roll vs. throw; blue), outcome (hit vs. miss; red), or a scenario × outcome interaction (yellow) in a sliding-window two-way ANOVA. Activity was aligned to contact (t = 0). **(H)** Percentage of physics-preferring and control-preferring units showing precontact tuning for outcome, scenario, and outcome × scenario interaction. Asterisk indicates *P* < 0.05; n.s., not significant. **(I)** Variance explained (η²) by outcome (top) and scenario (bottom) across the precontact period for physics-preferring and control-preferring units. Shaded regions indicate SEM; triangles indicate pointwise differences between populations (two-sided permutation tests, *P* < 0.05).

Population-averaged firing rates were higher for hit than miss trials in both scenarios, with the difference emerging 400 ms before contact (roll: 2.97 vs. 1.71 spikes/s; throw: 1.75 vs. 0.79 spikes/s; Bonferroni-corrected across time windows; Fig. 3B). Hit responses peaked at a median of 300 ms across units before contact in both scenarios, with no significant difference in time of peak firing relative to contact (Fig. 3C). The proportions of significantly responsive units were 29.4% and 27.3% for roll-hit and throw-hit trials, versus 24.4% and 17.7% for roll-miss and throw-miss trials (Fig. 3D). Responses were predominantly excitatory across all conditions (roll-hit: 73.2%; roll-miss: 66.2%; throw-hit: 67.2%; throw-miss: 63.6%; fig. S6B).

Precontact outcome selectivity was quantified using d’, with positive values indicating greater responses to hit than miss trials. The d′ distributions were significantly shifted above zero in both scenarios (median: 0.09 for roll, 0.06 for throw), with no significant difference between them (Fig. 3E). Outcome selectivity was also significantly correlated across scenarios (Spearman’s ρ = 0.14, *P* < 0.05; Fig. 3F), increasing to ρ = 0.36 (IQR [0.34, 0.38]) after correcting for within-scenario measurement reliability (see Materials and Methods).

The prevalence and temporal evolution of scenario and outcome tuning were further characterized using a two-way ANOVA across time (Fig. 3G). Scenario selectivity was most prevalent, peaking at 15.8% of units 200 ms before contact, followed by outcome selectivity (8.5% at the same time point). Scenario × outcome interactions were less prevalent, peaking at 3.6% at contact.

We next asked whether precontact outcome tuning was strongest among units showing consistent physics preference across the tower and video tasks (Fig. 3, H and I). Because the contact-prediction experiment was conducted during a subset of the sessions used for the physics analyses, we first confirmed that units recorded across all three tasks reproduced the main cross-task physics effects (fig. S7A). We then classified units as physics-preferring (d′ > 0 in both the tower and video tasks; n = 369) or control-preferring (d′ < 0 in both tasks; n = 76) and compared their precontact outcome tuning (see Materials and Methods). Outcome tuning was significantly more prevalent among physics-preferring units (12.2%, 45/369) than control-preferring units (1.3%, 1/76; Fisher’s exact test, odds ratio = 10.42, 95% CI [1.41, 76.78], *P* < 0.05). In contrast, the prevalence of scenario tuning (25.5% vs. 15.8%; odds ratio = 1.82, *P* > 0.05) and scenario × outcome interactions (2.4% vs. 3.9%; odds ratio = 0.61, *P* > 0.05) did not differ significantly between populations. Mean outcome-related explained variance was higher in physics-preferring units (η² = 0.02 vs. 0.01), with significant differences at individual precontact time bins (pointwise permutation tests, *P* < 0.05; Fig. 3I). Mean scenario-related explained variance was similar between populations (0.03 vs. 0.03), with no significant differences detected at any tested time bin.

Together, these results show that SPL responses differentiate upcoming collision outcomes across distinct physical scenarios, with outcome-related differences emerging before contact. Importantly, precontact outcome tuning was strongest among units that showed physics selectivity across both the tower and video tasks, linking intuitive physics representations to the prediction of upcoming physical outcomes.

### Population activity supports prediction of physical outcomes and generalization across scenarios

To assess whether collision outcomes could be decoded from SPL population activity and whether these representations generalized across scenarios, we performed population decoding analyses. Both scenario identity (roll vs. throw, pooling across outcomes) and collision outcome (hit vs. miss, pooling across scenarios) were robustly decoded, reaching peak accuracies of 96.2% and 94.2%, respectively (Fig. 4A). Significant decoding emerged well before contact, with scenario information appearing 700 ms and outcome information 600 ms before contact (first of at least two consecutive Bonferroni-corrected significant windows).

**Fig. 4.**
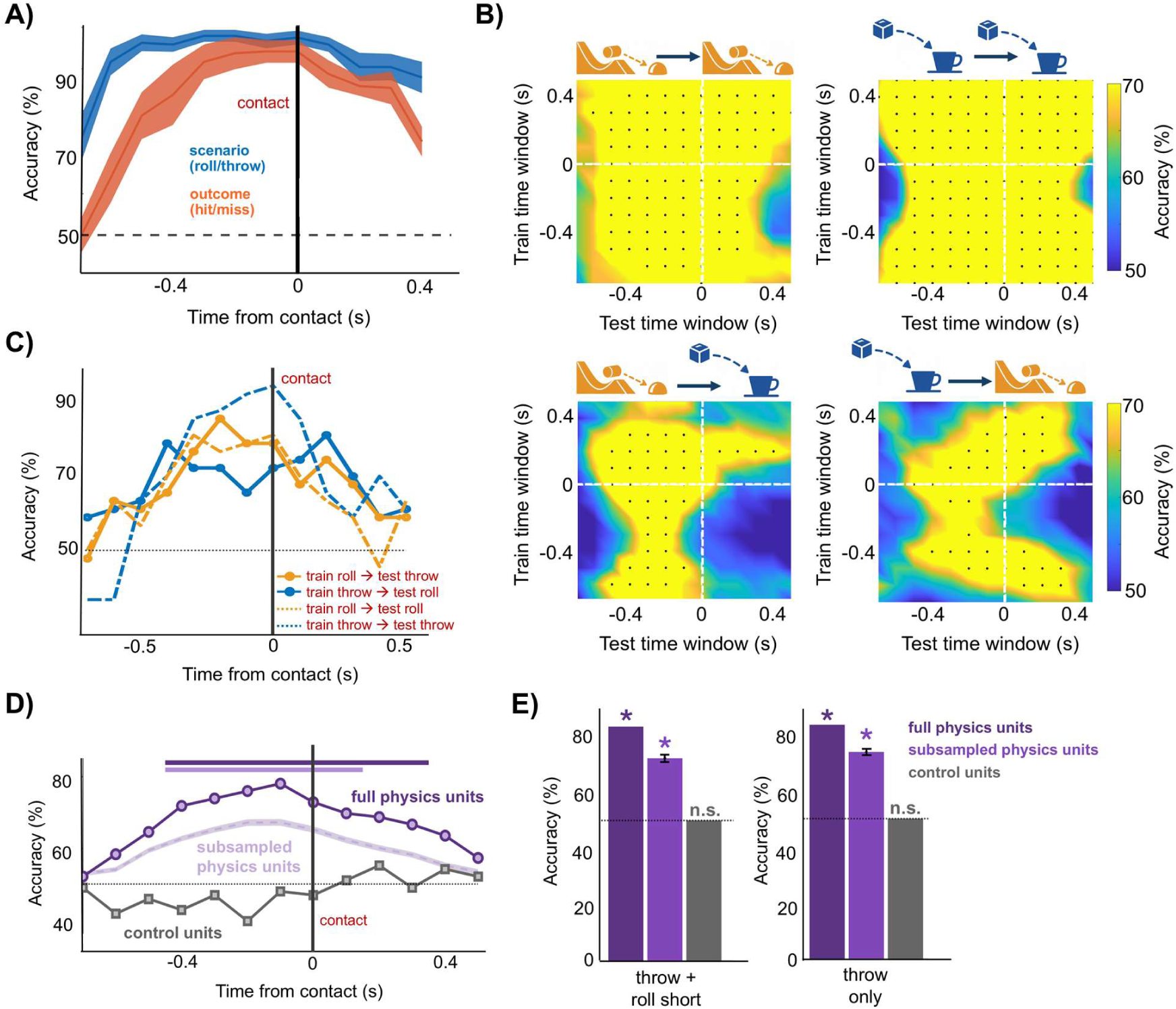
Population dynamics support prediction of physical outcomes before contact. **(A)** Time-resolved population decoding of scenario and outcome. Cross-validated LDA decoding accuracy for scenario (roll vs. throw; blue) and outcome (hit vs. miss; red) is shown as a function of time relative to contact (t = 0). Lines indicate mean decoding accuracy and shading indicates SEM. The horizontal dashed line indicates chance performance (50%). **(B)** Temporal generalization of outcome decoding within and across physical scenarios. Classifiers trained at each time window (y-axis) were tested at every time window (x-axis), either within the same scenario (roll → roll, top left; throw → throw, top right) or across scenarios (roll → throw, bottom left; throw → roll, bottom right). Color indicates hit-versus-miss decoding accuracy. Black dots indicate train–test time pairs with significant above-chance decoding based on permutation testing (Bonferroni-corrected across the full matrix, *P* < 0.05). Dashed white lines indicate contact (t = 0). **(C)** Time-resolved within- and cross-scenario outcome decoding. Diagonal decoding accuracies (matched train and test times) extracted from the temporal generalization matrices in (B) are shown for within-scenario decoding (dashed lines) and cross-scenario decoding (solid lines with markers). Orange indicates decoders trained on roll trials and blue indicates decoders trained on throw trials. The horizontal dotted line indicates chance performance (50%). **(D)** Cross-scenario outcome decoding from physics-preferring and control-preferring populations. Decoding accuracies at matched train and test time points (diagonal of the temporal generalization matrices) are shown for the full physics-preferring population (dark purple; n = 369), the control-preferring population (gray; n = 76), and physics-preferring units repeatedly subsampled to match the control-preferring population size (light purple; mean ± SEM). Horizontal bars indicate time windows with significant above-chance decoding for each population. The horizontal dotted line indicates chance performance (50%), and the vertical line marks contact (t = 0). Accuracy is averaged across roll-to-throw and throw-to-roll decoding; significant windows require both directions to meet the stated criterion. **(E)** Cross-latency generalization of outcome decoding in physics-preferring and control-preferring populations. Decoders were trained on either throw and short-latency roll trials (left) or throw trials alone (right) and tested on held-out long-latency roll trials using the precontact interval identified by the latency scan (−500 to −300 ms relative to contact). Accuracy is shown for the full physics-preferring population, the physics-preferring population subsampled to match the number of control-preferring units, and the control-preferring population. Error bars indicate SEM across 100 random neuron subsamples for the size-matched physics-preferring population. Asterisks indicate significant above-chance decoding (*P* < 0.05); n.s., not significant.

To test whether outcome information generalized across scenarios, we performed cross-temporal decoding of hit versus miss trials, training and testing at all pairs of time points within and across scenarios (Fig. 4B). Within-scenario decoding was robust, while cross-scenario decoding also emerged before contact in both directions. To quantify its onset, we examined decoding at matched train and test times (diagonal of the cross-temporal matrix; Fig. 4C). Using the same onset criterion, with permutation testing and Bonferroni correction across the full matrix, roll-to-throw decoding emerged 300 ms before contact and throw-to-roll decoding 400 ms before contact. Thus, SPL activity predicted collision outcomes before they occurred, generalizing across visually distinct physical scenarios.

To determine whether scenario-generalizable outcome information was preferentially carried by physics-preferring units, we repeated cross-scenario decoding separately for physics-preferring and control-preferring populations (Fig. 4D). Decoding was evaluated at matched train and test times, averaging both cross-scenario directions. Significant decoding in the physics-preferring population emerged 400 ms before contact (71.9% accuracy at onset). To control for population size, we repeated decoding with random subsamples of physics-preferring units matched in number to control-preferring units. Significant decoding again emerged 400 ms before contact (63.7% accuracy at onset), whereas the control-preferring population did not support significant cross-scenario decoding. When examined separately, all three populations supported significant precontact outcome decoding within individual scenarios (fig. S7B). In contrast, cross-scenario decoding was significant only in the physics-preferring populations. Direct comparisons further revealed significantly greater decoding accuracy in the size-matched physics-preferring population for throw-to-throw, throw-to-roll, and roll-to-throw decoding (one-sided normal-approximation tests, Bonferroni-corrected), but not for roll-to-roll decoding.

To assess whether cross-scenario predictive decoding could reflect differences in gaze rather than collision outcome, we compared precontact gaze position between hit and miss trials (see Materials and Methods). Horizontal gaze position, aligned to target side, differed significantly between outcomes in the throw scenario (*P* < 0.001) but not in the roll scenario (*P* > 0.05), while vertical gaze position did not differ significantly in either scenario (both *P* > 0.05). Because the outcome-related gaze difference was confined to the throw scenario, it cannot account for outcome decoding that generalized across scenarios.

Finally, to test whether outcome information was aligned to the upcoming collision rather than elapsed time from video onset, we examined cross-latency generalization in the different neuronal populations (Fig. 4E). Decoders were trained on either throw and short-latency roll trials or throw trials alone and tested on held-out long-latency roll trials, using a precontact window of −500 to −300 ms (see Materials and Methods and fig. S7, C and D). When trained on throw and short-latency roll trials, decoding generalized in both the full (83.3%, *P* < 0.05) and size-matched (72.1%, *P* < 0.05) physics-preferring populations, but not in control-preferring units (50.0%, *P* > 0.05). Training on throw trials alone yielded similar results (full: 83.3%; size-matched: 73.7%, both *P* < 0.05; control-preferring: 50.0%, *P* > 0.05). Thus, physics-preferring populations encoded collision outcomes relative to the upcoming contact event rather than stimulus onset.

Repeating all analyses using only well-isolated single units yielded consistent results for response latencies, outcome selectivity, ANOVA-based tuning, within- and cross-scenario decoding, and precontact decoding onset (fig. S6, C to L).

Our results show that SPL population activity contains predictive information about future physical outcomes hundreds of milliseconds before they occur, with several analyses converging on an onset approximately 400 ms before the event. Although outcome information was present more broadly across the neural population, only physics-preferring units carried an outcome representation that generalized across distinct physical scenarios. This generalizable representation was aligned to the upcoming contact event rather than to elapsed time from video onset, providing evidence that physics-preferring SPL populations encode predictions about future states of physical events. Together, these findings support a role for physics-preferring populations in the simulation function of a world model.

### Physics-preferring populations encode intended actions

Next, we tested whether SPL populations representing physical events also encode intended actions, as predicted if these units implement a physical world model. Participant RD performed an action-intention task involving either touching an object with his fist or pulling its knob with his thumb and index finger (Fig. 5A). Both actions elicited increased population activity, with firing rates significantly higher during touch than pull trials beginning 700 ms after the action cue (touch: 5.16; pull: 2.30 spikes/s across significant windows; Fig. 5B). Median response latency was 700 ms after the action cue, or 150 ms after the go cue (fig. S8A). Responses were predominantly excitatory during both touch and pull trials (67.1% and 78.7%, respectively; fig. S8B). Intended action identity was decoded with up to 95% accuracy (Fig. 5C), and cross-temporal decoding revealed stable representations across time (fig. S8C). These effects were also observed when analyses were restricted to well-isolated single units (fig. S9, A and B).

**Fig. 5.**
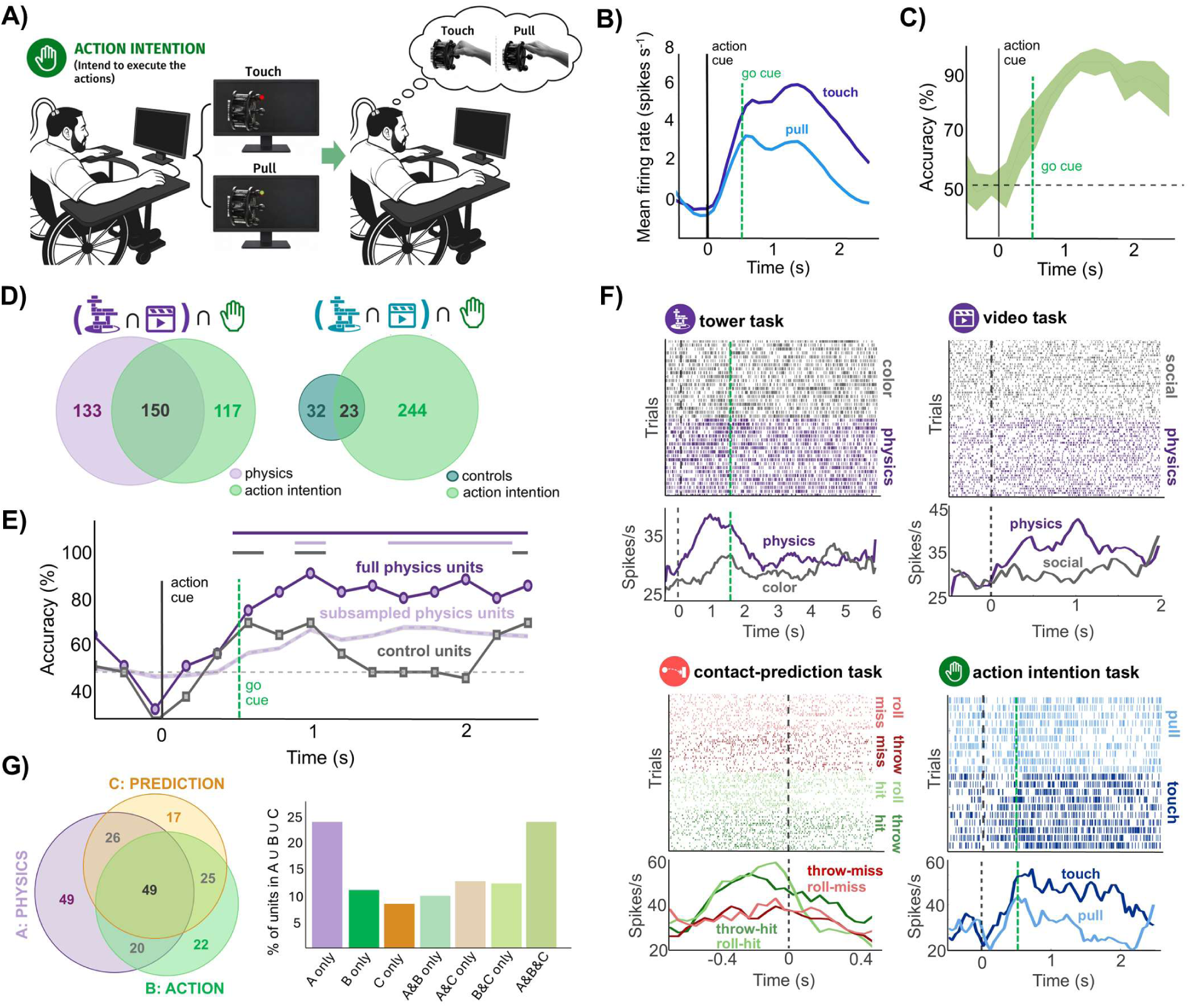
Shared neuronal populations link physical reasoning, prediction, and action planning. **(A)** Action-intention task. Participant RD viewed a cue instructing one of two actions, touch or pull, and, following a go cue, intended to execute the cued action. **(B)** Population-averaged firing rates during action intention, aligned to action-cue onset (t = 0), for touch (dark blue) and pull (light blue) trials. The dashed green line marks the go cue. **(C)** Time-resolved decoding of intended action (touch vs. pull). Cross-validated LDA decoding accuracy is shown relative to action-cue onset. The line indicates mean decoding accuracy and shading indicates SEM. The horizontal dashed line indicates chance performance (50%), and vertical lines mark action-cue onset and go cue. **(D)** Overlap between physics-preferring and action-responsive populations. Left, overlap between units consistently preferring physics over the respective control conditions in both the tower and video tasks (positive d′ in both tasks; purple) and units significantly modulated during action intention (green). Right, corresponding overlap for units consistently preferring the control conditions in both physics tasks (negative d′ in both tasks; teal) and action-responsive units. **(E)** Intended-action decoding within physics- and control-preferring populations. Touch-versus-pull decoding is shown using all physics-preferring units (*n* = 283; dark purple) and control-preferring units (*n* = 55; gray). To control for population size, physics-preferring units were repeatedly subsampled to *n* = 55 and decoding was repeated (light purple; mean ± SEM across subsamples). Horizontal bars indicate time windows with significant above-chance decoding and the horizontal dotted line indicates chance performance (50%). **(F)**Example SPL unit recorded across the four experimental paradigms. Raster plots (top) and corresponding trial-averaged firing rates (bottom) are shown for the tower task (physics and color), video task (physical and social), contact-prediction task (hit and miss outcomes for roll and throw scenarios), and action-intention task (touch and pull). Trials in raster plots are grouped by condition. Vertical dashed lines denote task events: tower-task prompt onset (black) and video onset (green); video onset in the video task and contact in the contact-prediction task; and action-cue onset (black) and go-cue onset (green) in the action-intention task. This unit exhibited greater firing during physics than control conditions in both the tower and video tasks, greater firing for hit than miss outcomes in both roll and throw scenarios, and was significantly modulated during action intention. **(G)** Overlap of physics-preferring, action-responsive, and contact-anticipatory units across the three sessions in which all four tasks were recorded from the same units (n = 262). Purple indicates physics-preferring units, defined by positive d′ in both the tower and naturalistic video tasks; green indicates action-responsive units, defined by significant modulation relative to baseline during action intention; and orange indicates contact-anticipatory units, defined by a significant modulation in firing rate before contact in at least one scenario-by-outcome condition. Numbers in the Venn diagram indicate unit counts for each mutually exclusive category. The bar plot shows the corresponding percentage of units in each category among units belonging to at least one of the three populations (n = 208).

We then tested whether intended action information was preferentially represented in the population of neurons identified as physics-preferring in the tower and video tasks. Because the action-intention task was performed in a subset of recording sessions, we first confirmed that these sessions reproduced the principal cross-task physics effects, including overlapping and correlated physics preferences (fig. S8D). Within this subset, units were classified as physics-preferring if they had positive d′ values in both tasks (n = 283) or control-preferring if they had negative d′ values in both (n = 55).

We examined the overlap between physics preference and action-related responsiveness (Fig. 5D). Of the 283 physics-preferring units, 150 (53.0%) showed significant modulation relative to baseline during action intention, accounting for 56.2% of all action-responsive units. Action-related responses were also observed in 23 of 55 control-preferring units (41.8%). Thus, although more than half of action-responsive units were physics-preferring, action-related activity was not exclusive to this population.

To determine whether physics-preferring units encoded intended action identity rather than merely responding during action intention, we first examined action tuning at the unit level. The proportion of action-tuned units did not differ significantly between physics-preferring (5.3%, 15/283) and control-preferring populations (3.6%, 2/55; Fisher’s exact test, odds ratio = 1.48, 95% CI [0.33, 6.68], P = 1). Mean variance in firing rate explained by intended action was also similar between populations (η² = 0.04 in both populations; fig. S8E).

At the population level, touch-versus-pull decoding was performed separately for physics-preferring and control-preferring units, with repeated subsampling of the former to match population sizes (n = 55; Fig. 5E). The full physics-preferring population supported robust decoding, reaching 95% accuracy at 900 ms. Size-matched decoding remained consistently above chance, peaking at 71.7% at 1750 ms, whereas decoding from control-preferring units was not consistently above chance. However, direct comparisons between size-matched populations revealed no significant differences at any time point. Thus, decoding of intended action identity remained consistently above chance in physics-preferring but not control-preferring populations, although decoding accuracy did not differ significantly between populations.

### Convergence of physics, prediction, and action representations in candidate world-model units

For the three recording sessions in which the tower, video, contact-prediction, and action-intention tasks were performed on the same units, we examined whether physics preference, collision anticipation, and action-related activity converged within individual units. Fig. 5F illustrates a candidate “world-model neuron” that showed physics preference across tasks, increased precontact activity for collisions in both throw and roll scenarios, and significant modulation during action intention.

Across the 262 units recorded in these sessions, 144 (55.0%) were physics-preferring, 117 (44.7%) were contact-anticipatory, and 116 (44.3%) were action-responsive (Fig. 5G). Physics preference was defined by positive d′ in both physics tasks, contact anticipation by significant precontact modulation in at least one scenario-by-outcome condition, and action responsiveness by significant modulation relative to baseline. Physics-preferring and contact-anticipatory populations overlapped significantly (75 units; odds ratio = 1.97, 95% CI [1.19, 3.24], *P* < 0.05), as did action-responsive and contact-anticipatory populations (74 units; odds ratio = 4.22, 95% CI [2.51, 7.10], *P* < 0.05). The overlap between physics-preferring and action-responsive populations was not significant (69 units; odds ratio = 1.39, 95% CI [0.85, 2.27], *P* > 0.05). Importantly, 49 units (18.7%) satisfied all three criteria, significantly exceeding chance overlap (permutation test, *P* < 0.05). This convergence persisted under stricter criteria for contact anticipation: requiring precontact modulation in both scenarios identified 22 units (8.4%), while additionally requiring modulation for the same outcome across scenarios identified 19 units (7.3%). Both overlaps remained significant (permutation tests, *P* < 0.05). Together, these findings identify a subset of SPL units exhibiting convergent representations of intuitive physics, anticipated physical events, and intended actions.

Qualitative inspection of the spatial distribution of physics-preferring, control-preferring, and action-responsive units revealed no obvious clustering across the electrode array, suggesting that these populations were intermingled within the recorded SPL region (fig. S9, C and D).

## Discussion

If the brain maintains an internal world model, a common neural substrate should integrate three functions essential for interacting with the physical world: representing the physical structure of the scene, predicting future states, and planning actions. Previous studies have implicated a frontoparietal network in intuitive physical inference and prediction (*14–17*), and posterior parietal cortex (PPC) in action planning (*11*, *13*, *18*), but whether these functions converge within the same neuronal populations remained unknown. Here, we show that the same SPL population, and indeed some of the same units, exhibited selectivity for physical information that generalized across tasks, predicted collision outcomes across scenarios, and encoded intended actions, providing neuronal-level evidence for a unified world model in human SPL.

This interpretation builds on a longstanding proposal that the nervous system constructs a “small-scale model” of external reality including possible actions, preserving the critical information relevant at a given moment rather than reproducing the world in full sensory detail (*1*). Modern computational world models similarly use compact latent representations to predict future states and evaluate actions (*8*, *19*, *20*). Li (*6*), building on earlier proposals in cognitive science and artificial intelligence (*21*), characterized world models for embodied agents in terms of three complementary capacities: a representation of the world’s state, its 3D geometry and physical properties; a simulator or dynamics model that predicts how the state will likely evolve in time; and a planner that uses these representations and predictive simulations to choose actions expected to lead to desired outcomes. Our three main findings map naturally onto these complementary components.

Physics selectivity during both explicit judgments and passive viewing may signal when a physical model should be constructed, whether prompted by the behavioral goal or the stimulus itself, consistent with our previous finding that SPL representations are gated by behavioral relevance (*22*). The stronger precontact outcome information in physics-preferring units links these representations to simulation, providing neuronal evidence for the prospective representations attributed to the cortical physics network (*17*). The same physics-preferring populations also carried information relevant to planning, although action tuning did not differ between populations and the overlap between physics preference and action responsiveness did not exceed chance (Fig. 5G; fig. S8E). This pattern is consistent with a distributed, mixed-selectivity representation in which intended-action information emerges from the joint activity of physics-preferring and non-preferring units.

Importantly, the selectivity of these responses cannot be explained simply by differential engagement or task difficulty (*23*, *24*). The color judgment control in the tower task was matched in difficulty to the stability judgment and engaged several processes associated with PPC, including approximate numerosity judgment (*25*), perceptual decision-making (*26*), and endogenous attention (*27*), yet elicited substantially weaker responses. Further, units were largely unresponsive to social videos (93.7% of units responded to physical videos versus 6.2% to social videos; Fig. 2B), which strongly engage other brain regions. Thus, although we observed mixed selectivity for physical and action information, these responses did not indiscriminately extend to other forms of cognitive demand (*23*, *24*), arguing against a generalized “everything everywhere” (*28*) representation.

Our results also provide a framework for interpreting established PPC computations as components of a world model. Studies in monkeys have shown that PPC represents targets in effector-centered coordinates (*18*), modulated by head and body position (*29*), and carries signals consistent with forward estimates of limb state (*30*). These computations could support transforming world representations into action plans and updating them as movements unfold.

The physical world model we identify in the SPL is part of a broader network engaged in intuitive physics, action planning, and tool use, including dorsal premotor cortex (PMd), supplementary motor area (SMA), and left supramarginal gyrus (SMG) (*14*, *31*). The connectivity, interactions, and division of labor between these regions in physical world modeling are important open questions for future research.

The world contains not only objects but also other agents whose actions shape how events unfold. Consistent with this, SPL responds during the observation of others’ actions (*32*), with similar neural responses to motion events caused by agents and by physical objects (*33*). The existence of neural populations that represent both observed and intended actions, originally described in monkeys (*34*), has recently been demonstrated in human SPL (*22*). As those findings were obtained from the same participants and arrays, they raise the possibility that shared representations of one’s own and others’ actions could form part of a broader world-model system (*31*). Consistent with this hypothesis, other regions of the physics network, including SMA, PMd, and SMG, are also recruited during action observation (*35*).

Our work provides an empirical and conceptual foundation, as well as the methods, for addressing several open questions. Does the same circuit simulate the consequences of possible actions to guide planning, and are these simulations computed through iterative, step-by-step updates? What prevents this circuit from engaging during social interactions, and might it be recruited when agents interact physically, as in fighting or dancing (*36*)? How does this circuit interact with broader brainwide networks? Answering these questions will help define the scope of this proposed world-model system.

Several limitations should be considered. Single-unit results were obtained from one participant, whereas the second contributed LFP recordings and performed only the physical reasoning tasks. Recordings were restricted to a small region of SPL, and the action task involved only two intended actions. The arrays also did not resolve cortical layers or cell types, leaving the underlying circuitry and interactions with the broader physics network and motor cortex unclear. Although our recordings cannot establish causality, parietal lesions have been associated with impaired novel tool use and mechanical problem solving, behaviors requiring the integration of physical reasoning and action planning (*37*). More fundamentally, physical representation, prediction, and action planning were tested separately, whereas in natural behavior these processes are tightly coupled. Whether and how the same neuronal populations integrate these functions within a single behavior remains unresolved.

We propose that these functions reflect different components of a common computation through which SPL models the state of the external world, anticipates its evolution, and plans actions within it. The foundation is now in place to unpack the computations and circuitry underlying the brain’s world model, with broad implications for human cognition, artificial intelligence, and the design of brain-computer interface devices aimed at restoring autonomy in people with paralysis.

## Materials and Methods

### Participants

Two right-handed male participants, JJ and RD (56 and 40 years old, respectively), were enrolled in a brain–computer interface clinical trial (ClinicalTrials.gov identifier: NCT01958086). The study was approved by the institutional review boards of the California Institute of Technology, Casa Colina Hospital and Centers for Healthcare, and the University of California, Los Angeles. Both participants provided informed consent before participation.

Participant JJ sustained a spinal cord injury at the C4–C5 level approximately 3 years before enrollment. He retained voluntary control of his eyes, head, and shoulders. Participant RD sustained a spinal cord injury at the C3–C4 level approximately 5 years before enrollment. He retained voluntary control of his eyes, head, and shoulders, as well as weak residual movements of his wrists and thumbs. Both participants were clinically stable during the study. Before implantation, functional magnetic resonance imaging confirmed task-related activation near the planned implant sites. Both participants were male; the influence of sex and gender on the results cannot be determined from this study and should be considered a limitation to generalizability.

### Electrophysiological Recordings

Participant JJ was implanted in 2018 with two 96-channel Utah microelectrode arrays targeting the left precentral gyrus and superior parietal lobule (SPL/JJ). Participant RD was implanted in 2023 with four 64-channel Utah arrays targeting two regions of the left precentral gyrus, the left superior parietal lobule (SPL/RD), and the left supramarginal gyrus. All arrays were 4 × 4 mm and had an interelectrode spacing of 400 µm (Blackrock Microsystems/Blackrock Neurotech, Salt Lake City, UT, USA).

The present study focused on recordings from the SPL arrays in both participants. Recordings from the precentral-gyrus arrays in both participants and the SMG array in participant RD were excluded because they did not exhibit reliable task-related modulation across recording sessions. Signal quality and unit isolation were comparable to other arrays. Given the anatomical and functional heterogeneity of SMG, the sampled subregion may not have overlapped with sites previously implicated in physical reasoning.

Neural signals were acquired using a 128-channel NeuroPort neural signal processor (Blackrock Neurotech). For participant RD, broadband signals were sampled at 30 kHz. Spiking activity was detected by applying a threshold of −3.5 times the root mean square of the high-pass-filtered signal. Recordings from participant RD included both single-unit and multiunit activity, as well as local field potentials (LFPs). LFPs were sampled continuously at 1 kHz.

For participant JJ, reliable spiking activity could no longer be isolated at the time of these experiments; therefore, only LFP signals were analyzed.

### Experimental setup

Experiments were conducted at Casa Colina Hospital and Centers for Healthcare for participant RD, and at home for participant JJ. In all sessions, participants remained seated in their motorized wheelchairs with their hands resting prone on a flat surface, in a well-lit room. A 30-inch LCD monitor was positioned directly in front of them. Stimulus presentation was controlled using the Psychophysics Toolbox for MATLAB (MathWorks).

#### Tower task

The tower task was adapted from Fischer et al. (*14*), using stimuli originally developed by Battaglia et al. (*7*). The stimuli consisted of 40 unique 6-s videos depicting unstable towers composed of blue, yellow, and white blocks. The towers were generated in Blender using rigid-body simulations, such that the blocks would collapse if gravity were allowed to take effect. Each tower was positioned at the center of a circular floor divided into red and green halves. During each video, the camera completed a 360° rotation around the stationary tower, allowing it to be viewed from multiple perspectives.

Participants performed either a physics judgment or a color judgment on each trial. Each trial began with a 500-ms intertrial interval, followed by a 1.5-s task cue. On physics trials, the cue asked, “Where will it fall?” Participants were instructed to predict whether more blocks would land on the red or green half of the floor. On color trials, the cue asked, “More blue or yellow?” Participants were instructed to determine whether the tower contained more blue or yellow blocks, disregarding the white blocks. The task cue was followed by the 6-s tower video, during which participants were instructed to maintain fixation at the center of the screen.

Following the video, participants reported their response using a saccade. On color trials, the response screen displayed the prompt “Color which had more blocks,” with a blue X on the left and a yellow X on the right. On physics trials, the response screen displayed the prompt “Color of the floor where more blocks will fall,” with a red X on the left and a green X on the right. Eye position was continuously monitored using a Tobii eye tracker. A response was registered when the participant fixated one of the two response targets for at least 500 ms. If no valid response was detected within 2 s, the trial ended and the experiment proceeded to the next trial.

The 40 unique tower videos were divided into two sets of 20. Within each session, every video was presented twice: once in the physics condition and once in the color condition. Each session therefore comprised 80 trials divided into two runs of 40 trials. Each run contained 20 unique videos, with every video presented once under each task condition. Trial order was randomized within each run and rerandomized for each recording session.

#### Video task

Participants passively viewed 72 naturalistic video clips divided equally between physical and social interaction categories. Each clip lasted 2 s and was preceded by a 500-ms intertrial interval. The 36 videos from each category were presented once per recording session, with all 72 videos presented in a fully randomized order.

Physical interaction videos depicted objects undergoing physical events (e.g., a door closing, a hammer driving a nail, falling dominoes, a spinning top, rolling dice). Social interaction videos depicted people engaged in social behaviors (e.g., laughing, crying, comforting one another, conversing, shaking hands).

Participants were instructed to maintain fixation on the screen while watching the videos and were permitted to blink naturally. Eye position was continuously monitored using a Tobii eye tracker; however, fixation was not enforced, and trials were not terminated or excluded based on gaze position. No behavioral response was required. Participants were not informed of the stimulus categories and were simply instructed to attend to the videos.

#### Contact prediction task

Participant RD passively viewed a set of 48 computer-generated 1.5-s video clips depicting interactions between a moving agent object and a stationary patient object, adapted from Pramod et al. (*17*). The stimuli followed a 2×2×6×2 factorial design, varying independently in scenario (roll or throw), outcome (hit or miss), background scene (six levels: three indoor and three outdoor), and direction of the agent object’s trajectory (leftward or rightward). This design yielded 48 unique videos, with 12 videos for each scenario-by-outcome combination. In the roll scenario, a cylinder rolled toward a stationary bowl. In the throw scenario, a cube moved through the air toward a stationary mug. All videos were created and rendered in Blender. The participant was instructed to maintain fixation while passively viewing the videos, and no behavioral response was required. Eye position was continuously monitored using a Tobii eye tracker; however, fixation was not enforced, and trials were neither terminated nor excluded based on gaze position. Each video was preceded by a 500-ms intertrial interval. Participant RD completed one run per recording session, during which each of the 48 videos was presented twice, resulting in 96 trials per session.

#### Action intention task

To compare neural responses during physical reasoning with those elicited during action planning, participant RD completed an action-intention task. Each trial lasted 2.5 s and began with the presentation of an object viewed from the side. A colored light on the object’s knob indicated which of two right-hand actions RD should intend and attempt: red cued touching the knob with a closed fist, whereas green cued pulling the knob with the thumb and index finger. The cue remained illuminated for 550 ms and then dimmed (go cue). After the cue dimmed, the object remained visible, but no hand or action was shown. RD was instructed to intend and attempt the cued action with his right hand during the remaining 1.95 s of the trial. Because he retained partial motor function, these attempts sometimes resulted in overt movement. Due to his physical limitations, the resulting movements did not necessarily reproduce the complete kinematics of the intended actions. Eye position was continuously monitored using a Tobii eye tracker. Each session comprised one 40-trial run, with 20 repetitions of each action. Trial order was fully randomized within each session.

### Behavioral choice accuracy (tower task)

Each response was scored according to the ground-truth answer for the corresponding video: the side of the floor on which more blocks landed for physics trials, or the more numerous block color for color trials. Trials involving towers with equal numbers of blue and yellow blocks had no defined correct response and were excluded from the color-accuracy analysis. Overall accuracy and accuracy within each task condition were compared against chance performance (50%) using two-sided exact binomial tests. Exact 95% Clopper–Pearson confidence intervals were calculated using the MATLAB binofit function. Differences in accuracy between the physics and color conditions were assessed using Fisher’s exact test on a 2×2 contingency table containing the numbers of correct and incorrect responses in each condition.

### Spatial distribution of gaze

To visualize spatial gaze patterns, we computed two-dimensional heatmaps of gaze position separately for each experimental condition, combining data across all trials and recording sessions. Gaze coordinates were concatenated within each condition, and two-dimensional histograms were computed using a fixed 100 × 100-bin grid spanning the full range of observed horizontal and vertical gaze positions. The resulting gaze-density distributions were interpolated for visualization to generate continuous heatmaps reflecting the spatial distribution of gaze during each condition.

### Single Unit Activity (SUA) and Multiunit Activity (MUA)

Each detected waveform consisted of 48 samples (1.6 ms total), including 10 samples before threshold crossing and 38 samples after. Single-unit and multiunit activity were sorted using Gaussian mixture modeling applied to the first three principal components of the waveform shapes (*22*, *38*). In addition to spike assignments, the sorting procedure provided a quality factor ranging from 1 to 4, determined by (i) the percentage of interspike intervals shorter than 3 ms, (ii) the signal-to-noise ratio of the mean waveform, (iii) the projection distance between clusters, (iv) the modified coefficient of variation of the interspike intervals (CV2), and (v) the isolation distance of each cluster. Units with a quality factor of 1 or 2 were considered well-isolated, whereas those with a factor of 3 or 4 were classified as multiunit activity (*22*, *38*). Following spike sorting, spike trains were converted to firing rates using 100-ms non-overlapping bins. For each unit, the mean firing rate was calculated across all trials and time bins within the analysis period, and units with a mean firing rate of 1 Hz or less were excluded from further analysis. For the retained units, firing rates were baseline-corrected on a trial-by-trial basis by subtracting the mean firing rate during the 500-ms prestimulus period (−500 to 0 ms) from the firing rate in each poststimulus bin.

### Local Field Potential (LFP)

Line noise was suppressed using a combined spectral and spatial filtering approach that preserves underlying neural signals while attenuating non-neural components (*39*). The data were then high-pass filtered above 2 Hz using a zero-phase infinite impulse response (IIR) filter. Trials in which the broadband signal amplitude exceeded two standard deviations from the session mean were excluded from further analysis. High-gamma (60–120 Hz) power was estimated using Morlet wavelet convolution with a 7-cycle resolution (*40*). To minimize edge artifacts introduced by filtering and wavelet convolution, the first and last 100 ms of each trial were discarded. Power was normalized within each trial by dividing the time–frequency power at each frequency by the mean power at that frequency during the prestimulus baseline.

### Comparison of LFP high-gamma power between conditions

To test whether high-gamma power (60–120 Hz) differed between task conditions (physics vs. color in the tower task; physical vs. social videos in the video task), power was averaged across recording channels and compared between conditions using an unpaired two-sided t-test in consecutive 250-ms windows advanced in 100-ms increments. P-values were corrected across all tested windows using the Bonferroni method, and a significant difference was defined as the center of the first window in a run of at least three consecutive Bonferroni-significant windows. Mean high-gamma power reported for each condition reflects the average, across all Bonferroni-significant windows, of that window’s trial-averaged power.

### Data aggregation

Unless otherwise specified, data were aggregated across recording sessions before analysis. For participant RD, spiking data from the SPL array were concatenated across sessions along the unit dimension, thereby treating units recorded in different sessions as a pooled population. Primary analyses included both single-unit and multiunit activity and were repeated using only single-unit activity. For participant JJ, SPL LFP data were concatenated across sessions along the trial dimension while preserving the electrode dimension.

### Cross-session unit-matching analysis

To quantify the potential re-identification of individual neurons across recording sessions, we compared units recorded on the same electrode across all pairs of sessions from participant RD, following the cross-session unit-matching method described in Bougou et al. (*22*).

For each unit, we calculated its mean spike waveform and interspike-interval distribution. Mean waveforms were centered by subtracting their temporal mean and then scaled to unit norm. Waveform similarity between two units was quantified as the dot product of their normalized mean waveforms. Similarity between interspike-interval distributions was quantified using the two-sample Kolmogorov–Smirnov statistic, with smaller values indicating more similar distributions. For each pair of recording sessions, we evaluated all cross-session pairs of units recorded on the same electrode. Null distributions were generated separately for each session pair by randomly sampling 5,000 cross-session unit pairs recorded on different electrodes. For each same-electrode pair, an empirical *P* value for waveform similarity was calculated as the proportion of null pairs with an equal or greater waveform similarity. An empirical *P* value for interspike-interval similarity was calculated as the proportion of null pairs with an equal or smaller Kolmogorov– Smirnov statistic. The two empirical *P* values were combined using Fisher’s method. Unit pairs with a combined *P* < 0.01 were classified as putative cross-session matches. For each pair of sessions, the match rate was calculated as the percentage of same-electrode unit pairs classified as putative matches.

### Contact-time alignment in the contact prediction task

For analyses of the contact-prediction task, neural activity was aligned to the time of contact in each video. Contact latency varied across stimuli. In the throw scenario, contact occurred consistently 800 ms after video onset. In the roll scenario, contact latencies ranged from approximately 820 to 1,450 ms, depending on the trajectory depicted in each clip. Roll stimuli were further separated into short-latency clips, in which contact occurred approximately 800–900 ms after video onset, and long-latency clips, in which contact occurred 1,100–1,500 ms after video onset.

To account for this variability, trials were re-epoched relative to the contact time associated with each video rather than analyzed exclusively relative to video onset. Contact times were assigned using a stimulus-specific lookup table derived from the video design (table S4). Because no physical contact occurred on miss trials, each miss video was assigned the contact time of its corresponding hit video, which differed only in the position of the target object.

For each trial, firing rates were extracted from 700 ms before to 500 ms after the assigned contact time. Samples were selected from the original baseline-corrected firing-rate time series using nearest-bin assignment at the native 100-ms resolution. Samples falling outside the recorded trial interval were treated as missing. This procedure yielded a common contact-relative time axis, with *t* = 0 denoting actual contact on hit trials and the corresponding hypothetical contact time on miss trials, permitting comparisons across stimuli and scenarios at matched times relative to contact.

### Significant task-related modulation and response latency

Task-related modulation was assessed independently for each unit and experimental condition: physics and color trials in the tower task; physical and social videos in the passive-viewing task; touch and pull trials in the action-intention task; and roll–hit, roll–miss, throw–hit, and throw–miss trials in the contact-prediction task. As described above, firing rates were baseline-corrected separately for each trial before statistical analysis. At each time window, the resulting net firing rates were compared with zero across trials using a two-sided one-sample *t*-test. For the tower, video, and action-intention tasks, firing rates were evaluated using 200-ms sliding windows advanced in 100-ms increments. Statistical significance was Bonferroni-corrected across all tested windows. A unit was classified as significantly modulated in a given condition if it reached the corrected significance threshold in at least three consecutive windows. Response latency was defined as the center of the first window in the earliest sequence satisfying this criterion. For the contact-prediction task, firing rates were evaluated in nonoverlapping 100-ms bins, with Bonferroni correction across all tested bins. A unit was classified as significantly modulated in a given scenario-by-outcome condition if it reached the corrected significance threshold in at least two consecutive bins. For units significantly modulated during “hit” trials, time of peak firing relative to contact was defined as the center of the bin containing the maximum trial-averaged firing rate during the precontact interval. Peak-response latencies were compared between the throw and roll scenarios using a two-sided Wilcoxon rank-sum test.

### Identification of action-responsive units

For analyses of population overlap, trials from the touch and pull conditions were pooled. For each unit, trial-wise baseline-corrected firing rates were compared with zero using two-sided one-sample t-tests in 200-ms windows advanced in 100-ms increments. P values were adjusted across time windows using the Benjamini–Hochberg false-discovery-rate procedure (q < 0.05). Units were classified as action-responsive if at least three consecutive windows reached significance.

### Comparison of firing rates between conditions

To test whether neural activity differed directly between conditions, trial-wise firing rates were first averaged across all units showing significant modulation in at least one condition (as defined above), yielding one population-averaged value per trial at each time point. These trial-level values were compared between conditions in consecutive 200-ms windows advanced in 100-ms increments using an unpaired two-sided t-test. P-values were corrected across all tested windows using the Bonferroni method, and a significant difference was defined as the center of the first window in a run of at least three consecutive Bonferroni-significant windows. Mean firing rates reported for each condition reflect the average, across all Bonferroni-significant windows, of that window’s trial-averaged population response.

### Excitatory and suppressive responses

For each experimental condition, all recorded units were separated according to the direction of their response relative to baseline. For each unit, the baseline-corrected firing rate was averaged across the task window within each trial and then across trials of the same condition. Units with a positive mean response were categorized as excited, whereas units with a negative mean response were categorized as suppressed. This analysis was performed separately for physics and color trials in the tower task; physical and social videos in the passive-viewing task; touch and pull trials in the action-intention task; and throw–hit, throw–miss, roll–hit, and roll–miss trials in the contact-prediction task. For each condition, the numbers of excited and suppressed units were expressed as percentages of all recorded units and displayed using diverging bar plots.

### Selectivity index

For each unit, selectivity between experimental conditions was quantified using the sensitivity index *d*’:

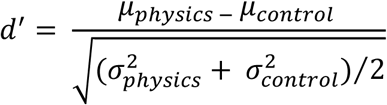

where *μ* and *σ*^2^ denote the mean and variance, respectively, of the trial-wise baseline-corrected firing rates within the relevant analysis window. For the tower task, *d*^’^ contrasted physics and color trials; for the video task, it contrasted physical and social videos; and for the contact-prediction task, it contrasted hit and miss trials separately within the throw and roll scenarios. Positive values indicated a preference for physics trials, physical videos, or hit outcomes, respectively, whereas negative values indicated a preference for the corresponding comparison condition.

### Cross-task overlap of physics-preferring units

A unit was defined as physics-preferring within a task when its *d*^’^ value was positive. For the video task, *d*^’^ was calculated over the full 2-s video presentation period, yielding a single set of physics-preferring units. For the tower task, *d*^’^ was calculated within a 2-s window advanced across the trial in 200-ms increments. At each window position, the overlap between the physics-preferring populations identified in the two tasks was quantified using the Jaccard index:

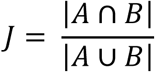

where *A* and *B* denote the sets of physics-preferring units in the tower and video tasks, respectively.

Statistical significance was assessed using a permutation test with 1,000 iterations. On each iteration, condition labels were independently permuted within each task, *d*^’^ values were recalculated, and the corresponding Jaccard index was obtained. The empirical *P* value at each tower task window was calculated as the proportion of permutations producing a Jaccard index equal to or greater than the observed value. P values were corrected across all tested windows using the Bonferroni method, and windows reaching this corrected threshold were considered significant. At the window with the largest significant Jaccard index, we additionally tested whether the distribution of d′ values across units was significantly different from zero in each task using two-sided Wilcoxon signed-rank tests. Differences in d′ magnitude between the tower and video tasks were assessed using a paired two-sided Wilcoxon signed-rank test. For visualization, the significant window with the largest significant observed Jaccard index was used to construct a proportional-area Venn diagram showing the numbers of units that were physics-preferring exclusively in the tower task, exclusively in the video task, or in both tasks.

### Cross-task correlation of condition preferences

To assess whether the magnitude and direction of condition preference were preserved across tasks, Spearman’s rank correlation was calculated between each unit’s tower task *d*^’^ and its video task *d*^’^. The passive-viewing *d*^’^ was calculated over the full 2-s video period, whereas tower task *d*^’^was recalculated within each 2-s window advanced in 200-ms increments. Statistical significance was assessed using 1,000 permutations. On each iteration, condition labels were independently permuted within each task, *d*^’^values were recalculated, and the cross-task Spearman correlation was recomputed at every tower task window. The empirical *P* value at each window was calculated as the proportion of permutations yielding a correlation coefficient equal to or greater than the observed coefficient. P values were corrected across all tested windows using the Bonferroni method. The 5th–95th percentile range of the permutation distribution was displayed as the null interval. Among the significant windows, the one with the largest significant correlation was selected as the best window, and a scatter plot of each unit’s video-task d′ against its tower task d′ (best window) was constructed, with units classified as consistently physics-preferring (*d*^’^ > 0 in both tasks), consistently color/social-preferring (*d*^’^ < 0 in both tasks), or sign-mismatched between tasks. The reported ρ and P correspond to the Spearman correlation and its asymptotic two-sided significance at this best window.

An analogous analysis was performed for the contact prediction task by correlating each unit’s outcome-selectivity *d*^’^ for hit versus miss trials in the throw scenario with its corresponding *d*^’^ in the roll scenario. Because this analysis was restricted to a single predefined precontact interval, the correlation was calculated once across the entire interval rather than across successive sliding windows. Statistical significance was assessed using the two-sided asymptotic test for Spearman’s rank correlation. In the corresponding scatter plot, units were categorized as consistently hit-preferring (*d*^’^ > 0 in both scenarios), consistently miss-preferring (*d*^’^ < 0 in both scenarios), or exhibiting opposite outcome preferences across scenarios.

### Within-task split-half reliability and disattenuation

To assess the reliability of unit-level d′ estimates and provide a reference for interpreting the cross-task correlation, we estimated within-task split-half reliability separately for each task. For each of 500 repetitions, trials were randomly divided into two halves within each condition, ensuring balanced condition representation across halves. d′ was recalculated independently for each half using the same analysis windows as for the corresponding correlation analysis: the window of strongest cross-task correlation for the tower task, the full 2-s video period for the video task, and the precontact interval for the throw and roll scenarios of the contact-prediction task. The resulting per-unit d′ values were correlated across halves using Spearman’s rank correlation. Split-half correlations were corrected to estimate full-sample reliability using the Spearman–Brown formula, *rSB* = 2*r*⁄(1 + *r*). The observed cross-task correlation was then corrected for attenuation using the estimated reliability of each task, 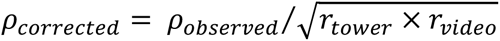. Disattenuated correlations are reported as the median and interquartile range across the 500 repetitions.

### Spatial distribution of selective units

To visualize the spatial distribution of physics-preferring, control-preferring, and action-responsive units across the array, each unit was assigned to its corresponding electrode on the physical 8 × 8 electrode layout. For each functional population, the number of units recorded on each electrode was displayed. Electrodes contributing at least one single unit to the corresponding population were marked separately. The resulting maps were used to qualitatively assess whether each population was spatially localized or distributed across the array.

### Significance of selectivity (d′ hit vs. miss) within and between scenarios

For each unit, outcome selectivity (*d*^’^, hit versus miss) was calculated over the entire precontact interval. Within each scenario, the distribution of *d*^’^ values across units was compared with zero using a two-sided Wilcoxon signed-rank test. Differences in outcome selectivity between the throw and roll scenarios were assessed using a paired, two-sided Wilcoxon signed-rank test. Pairing was performed by unit identity because each unit contributed one *d*^’^ value for each scenario.

### Control for intentional actions and visible body parts

To assess whether responses to physical-interaction videos were related to the presence of intentional actions or visible human body parts, two control analyses were performed using only physical-condition videos. Videos were manually categorized as depicting an intentional or non-intentional event (12 and 14 videos, respectively) and, independently, as containing or not containing a visible human body part (10 and 25 videos, respectively). Videos that could not be assigned unambiguously for a given comparison were excluded from that analysis. For each unit, *d*^’^ was calculated over the complete post-video-onset period for the intentional versus non-intentional and body-present versus body-absent contrasts. Positive values indicated greater responses to intentional-action or body-present videos, respectively. For each comparison, the distribution of *d*^’^ values across units was tested against zero using a two-sided Wilcoxon signed-rank test.

### Time-resolved two-way ANOVA in the contact-prediction task

To characterize the effects of outcome and scenario on neural activity during the contact prediction task, a two-way ANOVA was performed separately for each unit in consecutive, nonoverlapping 200-ms bins spanning the contact-aligned epoch. The model included outcome (hit or miss), scenario (throw or roll), and their interaction as factors, with trial-wise mean firing rate within each bin as the dependent variable. ANOVAs used Type III sums of squares. All trials were included in each model. At each time bin, *P* values were corrected across units using the Benjamini–Hochberg false-discovery-rate procedure (*q* < 0.05), separately for the main effects of outcome and scenario and for the outcome × scenario interaction. A unit was classified as exhibiting a main effect of outcome or scenario without interaction when the corresponding main effect reached significance and the interaction did not. Units with a significant interaction were considered separately. The proportions of units satisfying each criterion were calculated at each time bin relative to contact.

### Comparison of tuning between physics-preferring and control-preferring populations

To determine whether physics-preferring and control-preferring populations differed in their prevalence of outcome-, scenario-, and interaction-related tuning, we repeated the time-resolved two-way ANOVA described above, restricting the analysis to the precontact period. Units were classified as outcome-tuned if they exhibited a significant main effect of outcome without a significant outcome × scenario interaction in at least one precontact bin, and as scenario-tuned using the corresponding criterion for the main effect of scenario. Units exhibiting a significant outcome × scenario interaction in at least one precontact bin were classified as interaction-tuned. The proportions of units in each category were compared between the physics-preferring and control-preferring populations using Fisher’s exact tests on 2 × 2 contingency tables, with odds ratios, 95% confidence intervals, and *p* values reported for each comparison. As a complementary measure of tuning strength that did not depend on a significance threshold, we calculated η² separately for outcome and scenario for each unit in each precontact bin, as between-group sums of squares divided by total sums of squares. Differences in mean η² between physics-preferring and control-preferring units were assessed at each time bin using two-sided permutation tests with 500 random permutations of population labels.

### Comparison of action tuning between physics-preferring and control-preferring populations

To determine whether physics-preferring and control-preferring populations differed in intended-action tuning, a one-way ANOVA with intended action (touch versus pull) as the factor was performed separately for each unit in consecutive, nonoverlapping 100-ms bins spanning the post-cue period. ANOVAs used Type III sums of squares. At each time bin, *p* values were corrected across units using the Benjamini–Hochberg false-discovery-rate procedure (*q* < 0.05). Units were classified as action-tuned if they showed a significant effect of intended action in at least one post-cue bin. The proportions of action-tuned units were compared between the physics-preferring and control-preferring populations using Fisher’s exact test, with the odds ratio, 95% confidence interval, and *p* value reported. As a complementary measure of tuning strength independent of a significance threshold, we calculated η² for intended action for each unit in each post-cue bin, as between-group sums of squares divided by total sums of squares. Differences in mean η² between physics-preferring and control-preferring units were assessed at each time bin using two-sided permutation tests with 500 random permutations of population labels.

### Within-task population decoding

Time-resolved population decoding was performed using cross-validated linear discriminant analysis (LDA) run separately within each task: physics vs. color (tower task), physical vs. social (passive-viewing task), scenario (throw vs. roll) and outcome (hit vs. miss) (contact-prediction task, decoded separately), and touch vs. pull (action-intention task). For each cross-validation fold (10-fold), PCA was fit on the training trials only (pooled across all time bins), and the number of retained components was chosen to explain ≥95% of variance (capped at 20). A linear discriminant classifier was then trained and evaluated within matched, non-overlapping windows at each point in the trial, using the same time window for training and testing: 200-ms windows for the tower, passive-viewing, and action-intention tasks, and 100-ms windows for the contact-prediction task. Chance level was 50% (two classes in every case). Time was aligned to prompt onset for the tower task; to video onset for the passive-viewing task; to action-cue onset for the action-intention task; and to contact (via the contact-time realignment described above) for the contact-prediction task.

### Control for suppressed responses in video-task decoding

To determine whether decoding accuracy in the video task was driven by units showing suppressed (below-baseline) responses to social videos, units were classified as excited or suppressed separately for each condition, as described above (Excitatory and suppressive responses), based on the sign of their mean baseline-corrected firing rate across the full post-onset period. Physics-versus-social decoding was then repeated after excluding all units classified as suppressed during social videos, using the same cross-validated LDA procedure described above (Within-task population decoding).

### Cross-session robustness of population decoding

To determine whether decoding from the pooled cross-session pseudopopulation was disproportionately influenced by an individual recording session, two complementary control analyses were performed separately for the tower and video tasks. Both analyses used each session’s trial-wise mean firing rates over the corresponding post stimulus analysis window. First, condition identity was decoded independently within each recording session using only the units and trials recorded during that session. Physics versus color trials in the tower task and physical versus social trials in the video task were decoded using fivefold cross-validated LDA. This analysis assessed the consistency of condition information across individual recording sessions. Second, decoding was repeated using a leave-one-session-out pseudopopulation analysis. For each condition, pseudotrials were constructed by independently sampling, with replacement, one trial-wise response from each unit’s corresponding condition-specific response distribution. Thirty-six pseudotrials were generated per condition. This procedure was repeated across 20 independent resamples. Fivefold cross-validated LDA was applied to the complete pseudopopulation and to pseudopopulations generated after excluding all units from each recording session in turn. Decoding accuracy was averaged across resamples and compared between the complete and leave-one-session-out pseudopopulations to assess the contribution of individual sessions to the pooled decoding results. Pseudo-linear discriminant analysis was used when the number of units equaled or exceeded the number of trials in the smaller condition.

### Temporal cross-decoding

To assess whether each condition-identity code was temporally stable or reflected a sequence of transient states, the same classifier architecture was extended to a full temporal generalization analysis, performed separately within each task and for each decoded variable described above, using 10-fold cross-validation. For each cross-validation fold, PCA was fit once to the training trials pooled across all time bins, and the number of retained components was chosen to explain ≥ 95% of variance (capped at 20); the same projection was applied across all time windows. The LDA classifier was fit separately at each training window and evaluated at every test window: 200-ms windows for the tower, passive-viewing, and action-intention tasks, and 100-ms windows for the contact-prediction task, matching the corresponding matched-window decoding analysis. Held-out predictions were pooled across folds such that each trial contributed exactly one prediction for each train-time-by-test-time window pair. This yielded a train-time × test-time decoding-accuracy matrix for each task and decoded variable. Statistical significance at each matrix element was assessed using 1,000 permutations in which condition labels were shuffled relative to the pooled held-out predictions. P values were Bonferroni-corrected across all elements of the matrix. Decoding onset was defined as the center of the first of at least two consecutive time windows showing significant above-chance accuracy. For cross-scenario temporal generalization, significance was assessed after Bonferroni correction across all elements of the train-time × test-time matrix, and onset was determined from its diagonal.

### Cross-task and cross-scenario decoding

Cross-context generalization was evaluated between the tower and video tasks and between the throw and roll scenarios of the contact-prediction task. Tower–video decoding contrasted physics with color trials in the tower task and physical with social trials in the video task. Throw– roll decoding contrasted hit with miss trials. Decoding was performed bidirectionally (tower-to-video and video-to-tower; throw-to-roll and roll-to-throw) at every combination of training and test times.

For each decoding direction, activity was z-scored independently for each unit using the mean and standard deviation calculated from the complete training dataset; the same parameters were applied to the test dataset. PCA was fit to the normalized training data after pooling observations across time bins. The number of retained components was selected by fivefold cross-validation within the training dataset from 10 candidate dimensionalities evenly spaced between 2 and the maximum dimensionality. This maximum was set separately for each decoding type by inspecting cross-validated accuracy as a function of component count and choosing the smallest count beyond which additional components yielded only marginal further improvement (25 components for tower–video decoding; 15 components for throw–roll decoding). Because the training and test datasets were independent, no additional cross-validation was applied across contexts. A pseudo-linear discriminant classifier was fit at each 100-ms training bin and evaluated at every 100-ms test bin, yielding a train-time × test-time accuracy matrix. Chance accuracy was 50%.

For tower–video decoding, statistical significance was assessed using 1,000 permutations of the test labels relative to the fixed classifier predictions. Within each permutation, the same shuffled labels were applied across all train–test time pairs, and the maximum decoding accuracy across the matrix was retained. Each observed accuracy was compared with this maximum-statistic null distribution. Family-wise-error-corrected p values were calculated as (b + 1)/(1,000 + 1), where b was the number of permutation maxima equal to or greater than the observed accuracy. Matrix elements with corrected *P* < 0.05 were considered significant. For throw–roll decoding, significance was assessed with Bonferroni correction across all elements of the train-time × test-time matrix, as described above (Temporal cross-decoding).

Throw and roll trials were analyzed on the common contact-aligned time axis described above. Tower and video trials retained their respective task-relative time axes. For the contact-prediction task, within-scenario decoding was additionally performed as a reference using fivefold cross-validation, with folds grouped by video identity to prevent different repetitions of the same video from occurring in both the training and test sets. Out-of-fold predictions were pooled across folds. The matched-time within- and cross-scenario accuracies constitute the diagonal comparisons. Within-task tower and video decoding was obtained from the temporal generalization analysis described above and was not recomputed within the cross-task decoding pipeline.

### Eye position controls

To test whether gaze behavior could account for predictive decoding, we compared precontact eye position between hit and miss trials separately for the throw and roll scenarios. For each trial, horizontal and vertical gaze position were averaged over the 700-ms window preceding contact. Because the collision target appeared on the left or right side of the screen depending on the clip, horizontal gaze position was folded according to each clip’s target location, with positive values indicating gaze on the target side of the screen. This allowed pooling across clips without cancellation of target-related differences across left and right target locations. Vertical gaze position was analyzed without folding. Hit and miss trials (n = 192 each, pooled across sessions) were compared using two-sided independent-samples *t*-tests for each axis within each scenario, with a Bonferroni-corrected significance threshold of α = 0.0125 across the four comparisons. Horizontal gaze position differed between hit and miss trials in the throw scenario (hit, +1.87°; miss, −0.50°; *t*(382) = 7.91, *P* = 2.8 × 10⁻¹⁴) but not in the roll scenario (hit, +1.46°; miss, +1.13°; *t*(382) = 1.11, *P* = 0.27). Vertical gaze position did not differ in either scenario (throw, *t*(382) = 0.39, *P* = 0.70; roll, *t*(382) = −0.70, *P* = 0.49).

### Cross-latency generalization in the contact-prediction task

Cross-latency generalization was used to dissociate contact-relative outcome information from activity related to elapsed time since video onset. Contact occurred 800–900 ms after video onset in throw and short-latency roll trials (roll-short), but 1,100–1,500 ms after video onset in long-latency roll trials (roll-long). Thus, a hit–miss representation aligned to elapsed time from video onset would occur at different contact-relative times across the two latency groups, whereas a representation aligned to contact should generalize at matched times relative to contact. To identify the precontact interval for this analysis, we first performed a latency scan. Classifiers were trained using either the combined throw and roll-short trials (n = 84) or the throw trials alone (n = 48) and tested on the held-out roll-long trials. Using the same normalization, PCA, and pseudo-linear discriminant procedure described below, decoding was evaluated in 200-ms windows spaced 100 ms apart, spanning from 700 ms before contact to contact. Significance at each window was assessed using the trial-level permutation test described below (2,000 permutations), with Bonferroni correction across all tested windows applied separately for each training set. We identified the earliest window at which both training sets (throw alone and throw combined with roll-short) independently reached significance and used its center time to define a 200-ms precontact interval (±100 ms) for the primary cross-latency generalization analysis.

Given the limited number of roll-long trials (n = 12), these trials were reserved exclusively for testing. Classifiers were trained on activity within the identified precontact interval using either the combined throw and roll-short trials (n = 84) or the throw trials alone (n = 48). The former tested generalization across contact latency, whereas the latter tested generalization across both scenario and contact latency. PCA dimensionality was selected by fivefold cross-validation within each training dataset. A pseudo-linear discriminant classifier was then fit to the complete training dataset and applied, without refitting, to the roll-long trials within the corresponding contact-relative interval. Significance was assessed using 2,000 permutations of the roll-long outcome labels. For visualization, the fitted classifier was also applied across the complete contact-relative time course of the roll-long trials. Pointwise null intervals were derived from permutations of the test labels; these time-resolved values were used for visualization only and were not subjected to separate inferential testing.

### Restricted-population decoding of contact outcomes

To determine whether cross-scenario outcome information was preferentially represented by units with consistent physics-related response preferences, matched-time hit-versus-miss decoding was repeated separately using the physics-preferring and control-preferring populations defined by the cross-task *d*^’^analysis above. Decoding was evaluated at each bin of the shared contact-relative time axis for four training–testing configurations: throw-to-throw, roll-to-roll, throw-to-roll and roll-to-throw. Activity was z-scored using parameters derived exclusively from the training data. PCA was fit to the normalized training data pooled across time bins, and the number of retained components, up to a maximum of 15, was selected by fivefold cross-validated classification accuracy within the training dataset. A pseudo-linear discriminant classifier was then fit independently at each time bin and evaluated at the corresponding test bin. Within-scenario decoding used fivefold group cross-validation, with repeated presentations of the same video assigned to the same fold; held-out predictions were pooled across folds. For cross-scenario decoding, the training and test trials were disjoint, and the classifier was fit to the complete training scenario and evaluated in the other scenario. Accuracy was calculated separately for both cross-scenario directions and averaged to obtain a bidirectional summary. Statistical significance at each bin was assessed using 1,000 permutations of the test labels relative to the fixed classifier predictions; the bidirectional summary was considered significant only when both decoding directions independently reached *P* < 0.05. To control for differences in population size, the physics-preferring population was randomly subsampled without replacement to match the size of the control-preferring population, and decoding was repeated across 100 subsets using the same normalization, PCA and classification procedures. Accuracy was summarized as the mean ± SEM across subsets. A separate permutation distribution was not generated for each subset; instead, the 95th percentile of the control-preferring population’s permutation distribution was used as the size-matched chance threshold at each time bin. For the bidirectional summary, onset was defined as the center of the first of at least two consecutive bins in which both cross-scenario decoding directions independently reached significance at *P* < 0.05. At each time bin and for each training– testing configuration, the control-preferring population’s decoding accuracy was compared with the distribution of accuracies from 100 size-matched random subsets of physics-preferring units using a one-sided normal-approximation test. P values were Bonferroni-corrected across all tested time bins.

### Restricted-population cross-latency generalization in the contact-prediction task

The cross-latency generalization analysis described above was repeated separately for the physics-preferring and control-preferring populations defined by the cross-task d′ analysis, using the same precontact interval identified by the latency scan. As above, classifiers were trained using either the combined throw and roll-short trials (n = 84) or the throw trials alone (n = 48) and evaluated, without refitting, on the held-out roll-long trials. The same normalization, cross-validated PCA, and pseudo-linear discriminant procedure was used, with significance assessed using the trial-level permutation test (2,000 permutations) described above. To control for the larger number of physics-preferring units, physics-preferring units were randomly subsampled without replacement to match the size of the control-preferring population. The classification procedure was repeated across 100 independently drawn subsets using the same normalization, PCA, and classification procedure. Accuracy for the size-matched physics population was summarized as the mean ± SEM across subsets. Significance of the mean decoding accuracy was assessed using a matched permutation test in which the roll-long outcome labels were repeatedly shuffled and accuracy was averaged across the fixed predictions from all 100 subsampled classifiers for each permutation. This yielded a null distribution for the same across-subset mean accuracy statistic used to summarize the observed decoding performance.

### Restricted-population decoding of intended actions

To examine whether units exhibiting consistent physics-related preferences also encoded intended actions, touch-versus-pull decoding was repeated using unit populations defined by the cross-task *d*^’^ analysis described above. The physics-preferring population comprised units with positive *d*^’^ values in both the tower and video tasks, whereas the control-preferring population comprised units with negative *d*^’^ values in both tasks. Decoding was performed separately for the complete physics-preferring and control-preferring populations using nonoverlapping 200-ms windows. Within each fold of a fivefold cross-validation, PCA was fit once to the training data pooled across all time windows. The minimum number of components explaining at least 95% of the variance was retained, subject to a maximum of 20 components. The training and test data were projected into this component space, and a pseudo-linear discriminant classifier was fit and evaluated separately at each time window. Significance relative to chance was assessed separately for each population using 200 permutations of the trial labels. To control for the larger size of the physics-preferring population, units were randomly subsampled without replacement to match the size of the control-preferring population. Decoding was repeated for 100 independently drawn subsets using the same unit-wise normalization, fivefold cross-validation, PCA, and pseudo-linear LDA procedure. PCA dimensionality was selected separately for each subset and cross-validation fold using the same 95%-variance criterion and 20-component maximum. Decoding accuracy for the size-matched physics population was summarized as the mean ± SEM across subsamples. Because the two populations contained the same number of units, the 95th percentile of the permutation distribution obtained for the control-preferring population was used as the chance-reference threshold for the size-matched physics analysis; a separate permutation distribution was not generated for each subsample. Significant time windows were defined using the population-specific permutation distributions for the complete physics-preferring and control-preferring populations and the size-matched control-population threshold for the subsampled physics analysis. At each time window, the control-preferring population’s decoding accuracy was compared with the distribution of accuracies from 100 size-matched random subsets of physics-preferring units using a one-sided normal-approximation test. P values were Bonferroni-corrected across all tested time windows.

### Identification of “world-model” neurons

To identify units exhibiting convergent physics-, action-, and prediction-related responses, we analyzed the three sessions in which all four experimental tasks were recorded from the same units. Three functional populations were defined using criteria established in the corresponding analyses above. Physics-preferring units were defined as units with positive d′ in both the tower (physics > color) and naturalistic video (physical > social) tasks, using the cross-task time window identified by the Jaccard-index analysis described above. Action-responsive units were identified using the pooled-condition analysis described above under “Identification of action-responsive units.” Contact-anticipatory units were identified from firing rates in consecutive, nonoverlapping 100-ms bins spanning the precontact period. For each of the four scenario-by-outcome conditions (throw-hit, throw-miss, roll-hit, and roll-miss), firing rates were tested against zero using two-sided one-sample *t*-tests, with Bonferroni correction across time bins. A condition was considered to show significant anticipatory modulation when significance was observed in at least two consecutive bins. Because there is no established definition of a “world-model” neuron, we tested three progressively stricter criteria for defining contact-anticipatory units. Under the first criterion, units were required to show significant anticipatory modulation in at least one of the four scenario-by-outcome conditions. Under the second criterion, units were required to show significant anticipatory modulation in both scenarios, with at least one significant outcome condition within both the throw and roll scenarios. Under the third and most stringent criterion, units were required to show significant anticipatory modulation for the same outcome across both scenarios (i.e., throw-hit and roll-hit, or throw-miss and roll-miss). For each definition, pairwise associations between the three functional populations were assessed using Fisher’s exact tests on 2 × 2 contingency tables. To determine whether the three-way overlap exceeded that expected by chance, the physics-preferring and action-responsive populations were held fixed while unit identities for the contact-anticipatory population were randomly reassigned, preserving its observed population size. This procedure was repeated 2,000 times, and the number of resampled contact-anticipatory units falling within the intersection of the physics-preferring and action-responsive populations was calculated at each iteration. The empirical *p* value was defined as the proportion of permutations in which the resulting three-way overlap was equal to or greater than the observed overlap.

## Acknowledgments

We would like to express our gratitude to participants JJ and RD for making this research possible. We would like to thank Vivian Paulun and Thomas Brewitt for the stimuli used in the Video Task. During manuscript preparation, we used ChatGPT (OpenAI) for language editing and text refinement.

## Funding

National Eye Institute of the National Institutes of Health grant UG1EY032039 (RAA)

Tianqiao and Chrissy Chen Brain-Machine Interface Center at Caltech (RAA)

Swartz Foundation (RAA)

Boswell Foundation (RAA)

Fonds Wetenschappelijk Onderzoek (FWO) grant 1264926N (VB)

MIT Siegel Family Quest for Intelligence (NK)

Office of Naval Research MURI grant N00014-21-1-2801 (JBT)

Office of Naval Research MURI grant N00014-22-1-2740 (JBT)

Office of Naval Research MURI grant N00014-24-1-2603 (JBT)

## Author contributions

Conceptualization: VB, RTP, JBT, NK, RAA

Methodology: VB, RTP, JBT, NK, RAA

Investigation: VB

Formal analysis: VB

Resources: ERR, AB

Funding acquisition: VB, KP, RAA, JBT, NK

Project administration: KP, RAA

Supervision: NK, RAA

Writing – original draft: VB

Writing – review & editing: VB, RTP, ERR, AB, KP, JBT, NK, RAA

## Competing interests

Authors declare that they have no competing interests.

## Data, code, and materials availability

Data and analysis code will be deposited in CaltechDATA, with the repository DOI provided before publication.

## Notes

### Competing Interest Statement

The authors have declared no competing interest.

### Summary of Updates

Corrected the layout and formatting of figures and figure legends. No changes were made to the scientific content.

## References

1. K. J. W. Craik, The Nature of Explanation (Cambridge University Press, Cambridge, 1943).

2. P. N. Johnson-Laird, Mental Models: Towards a Cognitive Science of Language, Inference, and Consciousness (Harvard University Press, Cambridge, MA, 1983).

3. D. M. Wolpert, Z. Ghahramani, M. I. Jordan, An internal model for sensorimotor integration. Science 269, 1880–1882 (1995).

4. D. M. Wolpert, Z. Ghahramani, Computational principles of movement neuroscience. Nat. Neurosci. 3, 1212–1217 (2000).

5. Y. LeCun, A Path Towards Autonomous Machine Intelligence. (2022). https://openreview.net/forum?id=BZ5a1r-kVsf.

6. F. F. Li, A Functional Taxonomy of World Models (2026). https://drfeifei.substack.com/p/a-functional-taxonomy-of-world-models.

7. P. W. Battaglia, J. B. Hamrick, J. B. Tenenbaum, Simulation as an engine of physical scene understanding. Proc. Natl. Acad. Sci. U. S. A. 110, 18327–18332 (2013).

8. D. Ha, J. Schmidhuber, Recurrent World Models Facilitate Policy Evolution. Adv. Neural Inf. Process. Syst. 31 (2018).

9. M. A. Goodale, A. D. Milner, Separate visual pathways for perception and action. Trends Neurosci. 15, 20–25 (1992).

10. J. C. Culham, K. F. Valyear, Human parietal cortex in action. Curr. Opin. Neurobiol. 16, 205–212 (2006).

11. R. A. Andersen, C. A. Buneo, Intentional maps in posterior parietal cortex. Annu. Rev. Neurosci. 25, 189–220 (2002).

12. R. A. Andersen, H. Cui, Intention, Action Planning, and Decision Making in Parietal-Frontal Circuits. Neuron 63, 568–583 (2009).

13. T. Aflalo, S. Kellis, C. Klaes, B. Lee, Y. Shi, K. Pejsa, K. Shanfield, S. Hayes-Jackson, M. Aisen, C. Heck, C. Liu, R. A. Andersen, Decoding motor imagery from the posterior parietal cortex of a tetraplegic human. Science 348, 906–910 (2015).

14. J. Fischer, J. G. Mikhael, J. B. Tenenbaum, N. Kanwisher, Functional neuroanatomy of intuitive physical inference. Proc. Natl. Acad. Sci. U. S. A. 113, E5072–E5081 (2016).

15. S. E. Schwettmann, J. B. Tenenbaum, N. Kanwisher, Invariant representations of mass in the human brain. elife 8, e46619 (2019).

16. R. T. Pramod, M. A. Cohen, J. B. Tenenbaum, N. Kanwisher, Invariant representation of physical stability in the human brain. elife 11, e71736 (2022).

17. R. T. Pramod, E. Mieczkowski, C. X. Fang, J. B. Tenenbaum, N. Kanwisher, Decoding predicted future states from the brain’s “physics engine.” Sci. Adv. 11, eadr7429 (2025).

18. L. H. Snyder, A. P. Batista, R. A. Andersen, Coding of intention in the posterior parietal cortex. Nature 386, 167–170 (1997).

19. D. Hafner, T. Lillicrap, J. Ba, M. Norouzi, Dream to Control: Learning Behaviors by Latent Imagination. Int. Conf. Learn. Represent. (2019).

20. D. Hafner, J. Pasukonis, J. Ba, T. Lillicrap, Mastering diverse control tasks through world models. Nature 640, 647–653 (2025).

21. B. M. Lake, T. D. Ullman, J. B. Tenenbaum, S. J. Gershman, Building machines that learn and think like people. Behav. Brain Sci. 40, e253 (2017).

22. V. Bougou, J. Gamez, E. R. Rosario, C. Liu, K. Pejsa, A. Bari, R. A. Andersen, Hierarchical and context-dependent encoding of actions in human posterior parietal and motor cortex. Cell 189, 5304–5317.e7 (2026).

23. J. Duncan, The multiple-demand (MD) system of the primate brain: mental programs for intelligent behaviour. Trends Cogn. Sci. 14, 172–179 (2010).

24. E. Fedorenko, J. Duncan, N. Kanwisher, Broad domain generality in focal regions of frontal and parietal cortex. Proc. Natl. Acad. Sci. U. S. A. 110, 16616–16621 (2013).

25. A. Nieder, S. Dehaene, Representation of number in the brain. Annu. Rev. Neurosci. 32, 185–208 (2009).

26. J. I. Gold, M. N. Shadlen, The neural basis of decision making. Annu. Rev. Neurosci. 30, 535–574 (2007).

27. E. Wojciulik, N. Kanwisher, The generality of parietal involvement in visual attention. Neuron 23, 747–764 (1999).

28. M. C. Rosen, D. J. Freedman, How distributed is the brain-wide network that is recruited for cognition? Nat. Rev. Neurosci. 27, 138–150 (2026).

29. L. H. Snyder, K. L. Grieve, P. Brotchie, R. A. Andersen, Separate body- and world-referenced representations of visual space in parietal cortex. Nature 394, 887–891 (1998).

30. G. H. Mulliken, S. Musallam, R. A. Andersen, Forward estimation of movement state in posterior parietal cortex. Proc. Natl. Acad. Sci. U. S. A. 105, 8170–8177 (2008).

31. J. Fischer, B. Z. Mahon, What tool representation, intuitive physics, and action have in common: The Brain’s First-Person Physics Engine. Cogn. Neuropsychol. 38, 455–467 (2021).

32. J. Decety, J. Grèzes, Neural mechanisms subserving the perception of human actions. Trends Cogn. Sci. 3, 172–178 (1999).

33. S. Karakose-Akbiyik, O. Sussman, M. F. Wurm, A. Caramazza, The Role of Agentive and Physical Forces in the Neural Representation of Motion Events. J. Neurosci. 44, e1363232023 (2024).

34. G. di Pellegrino, L. Fadiga, L. Fogassi, V. Gallese, G. Rizzolatti, Understanding motor events: a neurophysiological study. Exp. Brain Res. 91, 176–180 (1992).

35. S. Caspers, K. Zilles, A. R. Laird, S. B. Eickhoff, ALE meta-analysis of action observation and imitation in the human brain. Neuroimage 50, 1148–1167 (2010).

36. S. Liu, S. Karakose-Akbiyik, J. Outa, M. J. Kim, How physical information is used to make sense of the psychological world. Nat. Rev. Psychol. 5, 59–73 (2026).

37. G. Goldenberg, J. Spatt, The neural basis of tool use. Brain 132, 1645–1655 (2009).

38. C. Y. Zhang, T. Aflalo, B. Revechkis, E. R. Rosario, D. Ouellette, N. Pouratian, R. A. Andersen, Partially Mixed Selectivity in Human Posterior Parietal Association Cortex. Neuron 95, 697–708.e4 (2017).

39. A. de Cheveigné, ZapLine: A simple and effective method to remove power line artifacts. Neuroimage 207 (2020).

40. R. Kronland-Martinet, J. Morlet, A. Grossmann, Analysis of Sound Patterns through Wavelet transforms. Int. J. Pattern Recognit. Artif. Intell. 01, 273–302 (1987).

