## Supplementary Material for "A Unified World Model in Human Posterior Parietal Cortex"

**Fig. S1. Stability of single-unit recordings across sessions.**

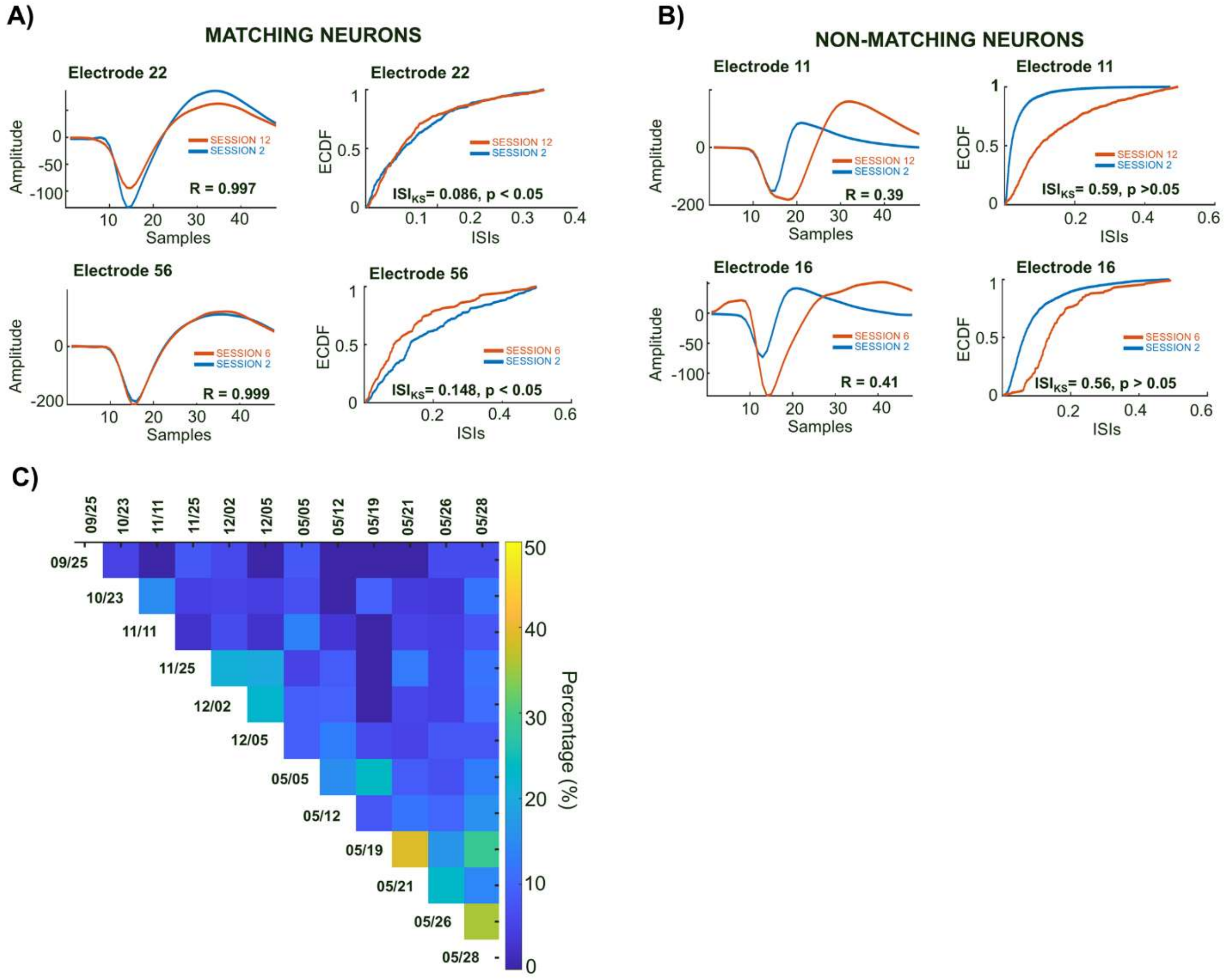


**(A)** Examples of matching neurons recorded from the same electrode across two sessions, as identified from the cross-session unit matching analysis. Left panels show average spike waveforms (48 samples, 1.6 ms) aligned to threshold crossing for two recording days (red and blue). Correlation coefficients (R) quantify waveform similarity across sessions. Right panels show empirical cumulative distribution functions (ECDFs) of inter-spike intervals (ISIs) for the same units across sessions; Kolmogorov–Smirnov statistics and p-values are indicated. High waveform correlations and overlapping ISI distributions demonstrate stable unit isolation across days. **(B)** Examples of non-matching neurons recorded from the same electrode across sessions. Waveform shapes differ substantially between sessions (lower R values), and ISI distributions show marked divergence, consistent with distinct units recorded on different days. **(C)** Percentage of same-electrode unit pairs classified as putative cross-session matches across pairs of recording dates. Each cell indicates the proportion of same-electrode unit pairs meeting the waveform similarity and ISI criteria for the corresponding session pair. Warmer colors indicate higher match rates.

**Fig. S2. Gaze, LFP and response-direction controls for physics selectivity in the tower and video tasks.**

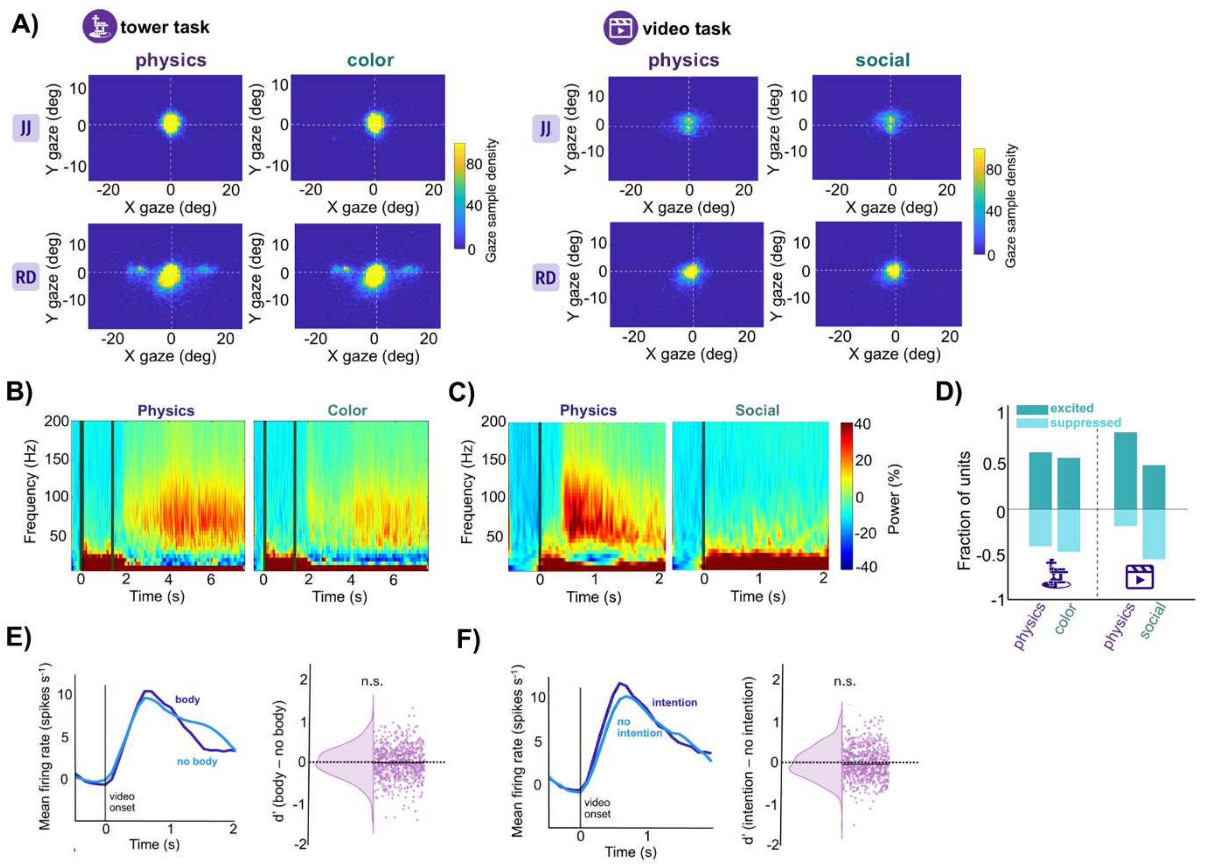


**(A)** Gaze distributions during the tower and video tasks. Heatmaps show the two-dimensional distribution of gaze position, in degrees of visual angle, aggregated across trials for JJ and RD and shown separately for each condition. Gaze remained predominantly centered across participants, tasks, and conditions. Panels B–F show data from participant RD. **(B and C)** Time-frequency representations of LFP power during the tower task (B) and video task (C), averaged across trials and recording channels. Physics conditions are shown on the left and the corresponding color or social control conditions on the right. Color indicates change in spectral power relative to baseline. **(D)** Direction of task-evoked responses across conditions. Bars show the fraction of all recorded units exhibiting increased (excited; dark teal) or decreased (suppressed; light teal) firing relative to baseline for the physics and color conditions of the tower task and the physical and social conditions of the video task. **(E)** Control for the presence of visible body parts in the physical clips of the video task. Left, population-averaged firing rates for physical videos containing versus not containing visible body parts. Right, distribution of unit-level d′ values comparing body versus no-body videos (median: -0.02). The distribution did not differ significantly from zero (n.s.). **(F)** Control for intentional versus non-intentional physical events in the physical clips of the video task. Left, population-averaged firing rates for physical videos depicting events involving versus not involving intentional actions. Right, distribution of unit-level d′ values comparing intentional versus non-intentional videos (median: -0.05). The distribution did not differ significantly from zero (n.s.).

**Fig. S3. Example SPL neurons showing physics selectivity across the tower and video tasks.**

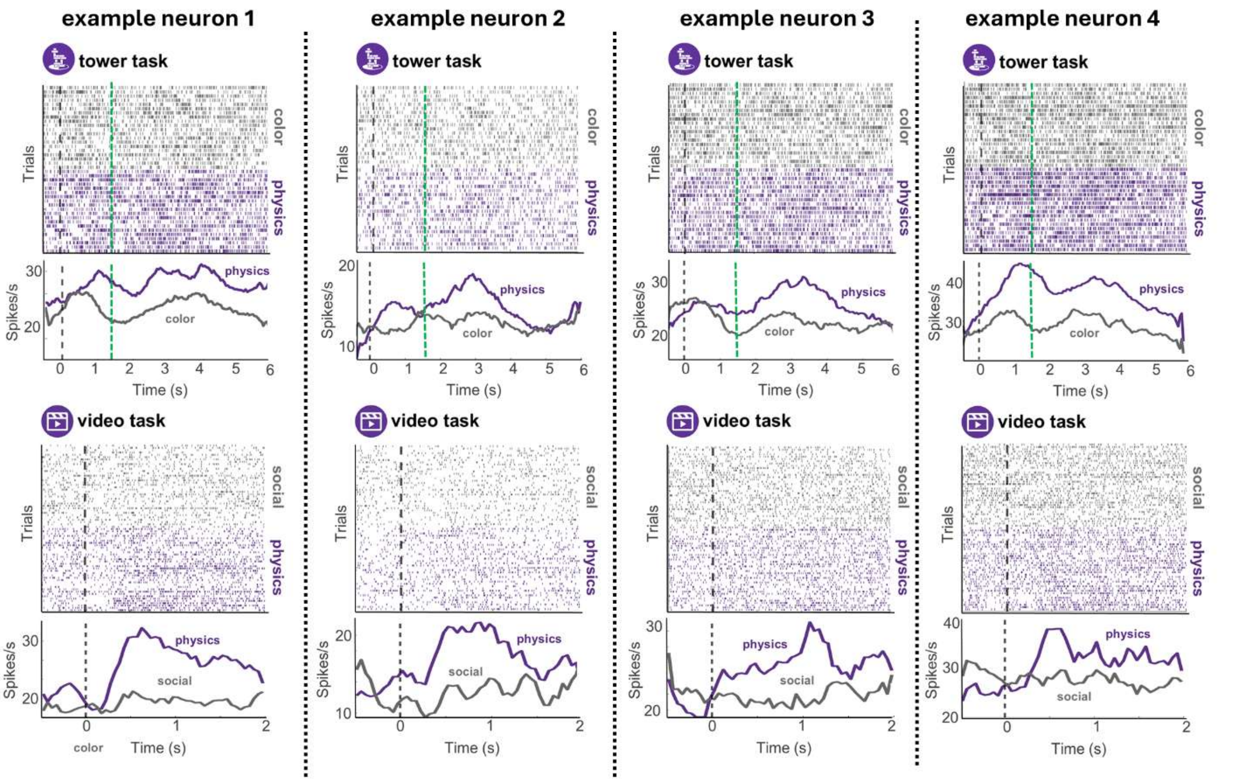


Four example SPL neurons showing physics selectivity across the tower and video tasks. For each unit, raster plots (top) and corresponding trial-averaged firing rates (bottom) are shown for the tower task (physics and color) and video task (physical and social). Trials in raster plots are grouped by condition. In the tower task, vertical dashed lines denote prompt onset (black) and movie onset (green); in the video task, the vertical dashed line denotes video onset. All four neurons showed greater responses to physics than control conditions in both tasks.

**Fig. S4. Population decoding controls: suppressive responses, temporal generalization, and session stability.**

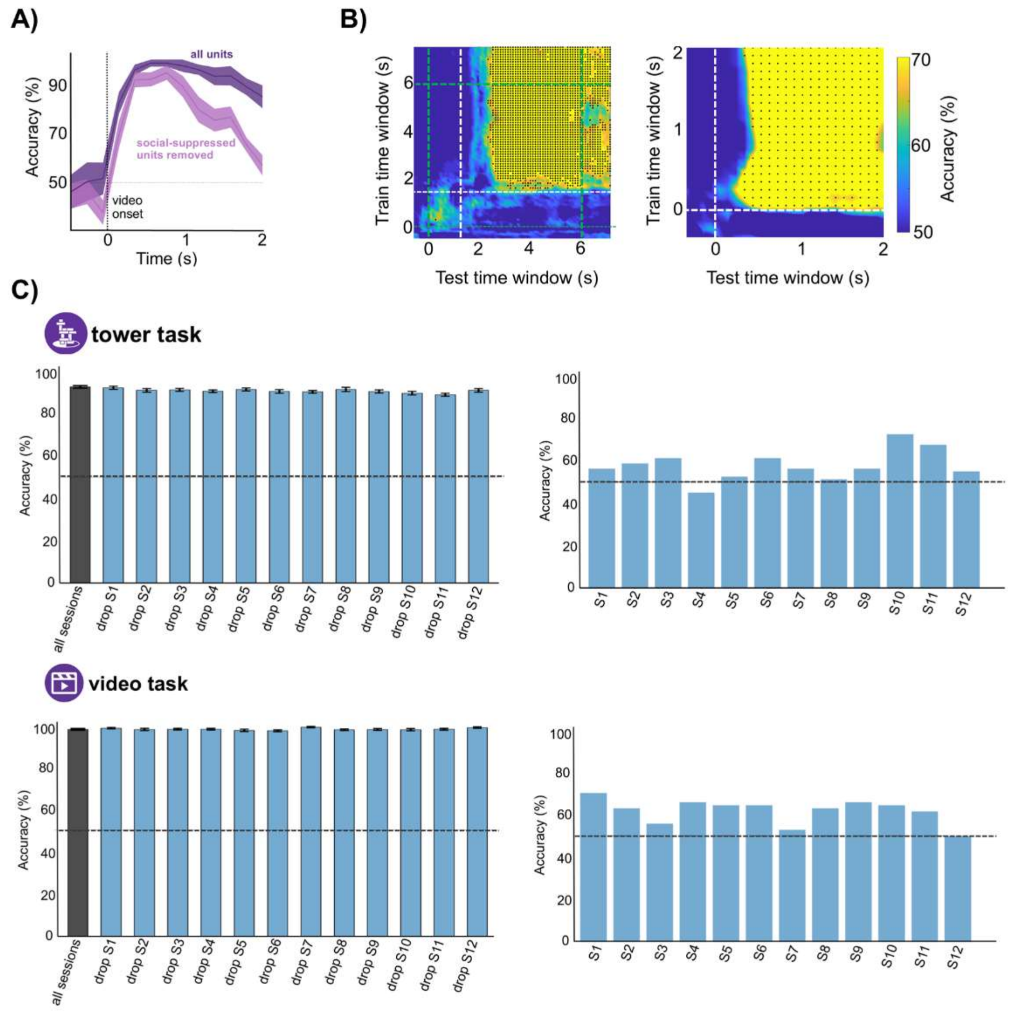


**(A)** Population decoding accuracy for discriminating physical from social videos using all units or excluding units showing suppression during social videos. Shaded regions indicate SEM across cross-validation folds. The dashed line indicates chance level (50%). **(B)** Temporal generalization of physics-versus-control decoding within the tower task (left) and video task (right). Classifiers trained at each time window (y-axis) were tested across all time windows (x-axis). Color indicates decoding accuracy; black dots indicate significant train–test pairs. Dashed lines mark task events. **(C)** Session-stability controls for physics decoding. Top row, tower task (physics versus color). Left, leave-one-session-out pseudopopulation decoding accuracy using all sessions (gray) or after excluding each session in turn (blue). Bars show mean accuracy across repetitions; error bars indicate SEM. Right, within-session decoding accuracy computed independently from simultaneously recorded units and single-trial responses in each session. Bottom row, same analyses for the video task (physical versus social). Dashed horizontal lines indicate chance performance (50%).

**Fig. S5. Cross-task physics representations are preserved in single-unit activity.**

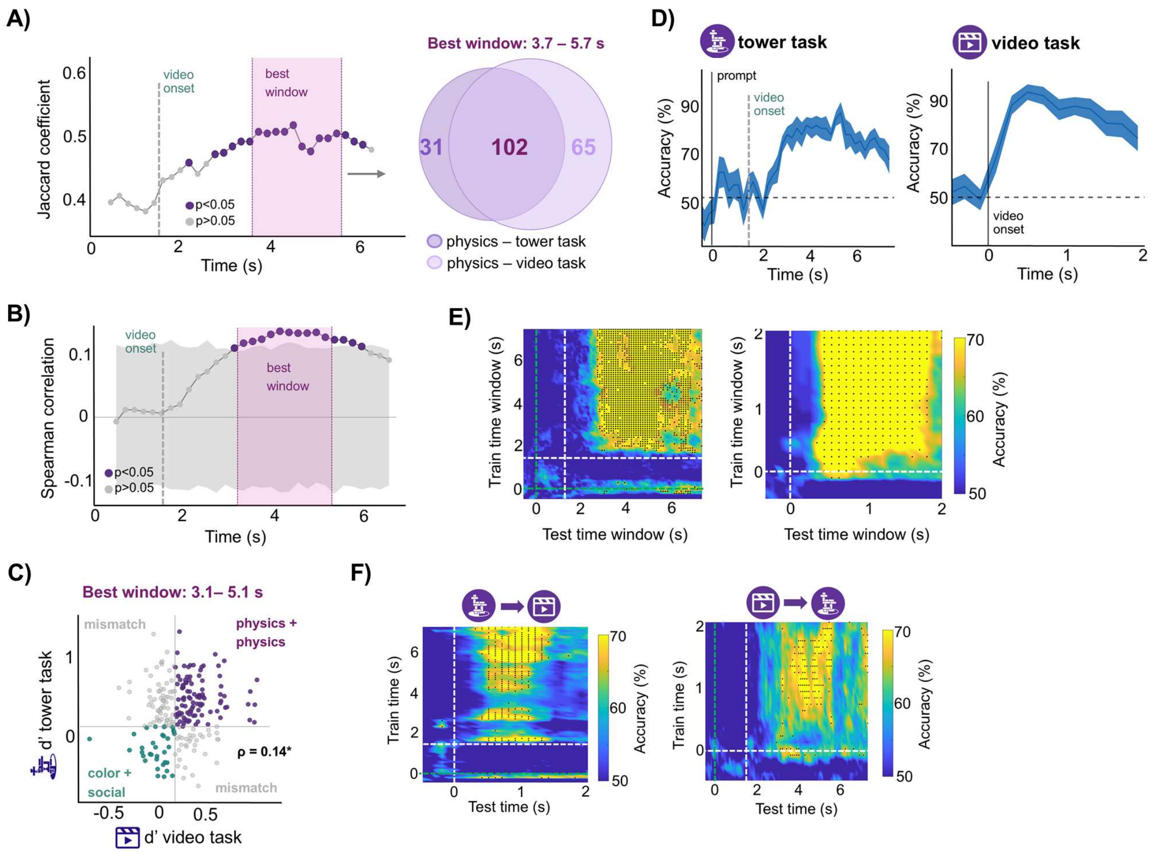


Analyses from Fig. 2 were repeated using only well-isolated single units (SUA). **(A)** Overlap of physics-selective single units across the tower and video tasks, quantified by the Jaccard coefficient across tower-task analysis windows (left) and at the best-overlap window (3.7–5.7 s; right). At the best window, 102 units were physics-selective in both tasks, with 31 selective only in the tower task and 65 only in the video task. **(B)** Spearman correlation between single-unit physics selectivity across tasks as a function of the tower-task analysis window. **(C)** Relationship between tower- and video-task physics selectivity within the best window (3.1–5.1 s). Physics selectivity was significantly correlated across tasks (Spearman $\rho = 0.14$, $P < 0.05$). **(D)** Time-resolved LDA decoding of physics versus control conditions within the tower and video tasks, reaching peak accuracies of 83.7% and 91.8%, respectively. **(E)** Temporal generalization of decoding within each task. **(F)** Cross-task temporal generalization of physics decoding, with classifiers trained in the tower task and tested across time in the video task (left), and trained in the video task and tested across time in the tower task (right).

**Fig. S6. Contact-outcome encoding and generalization are preserved in single-unit activity.**

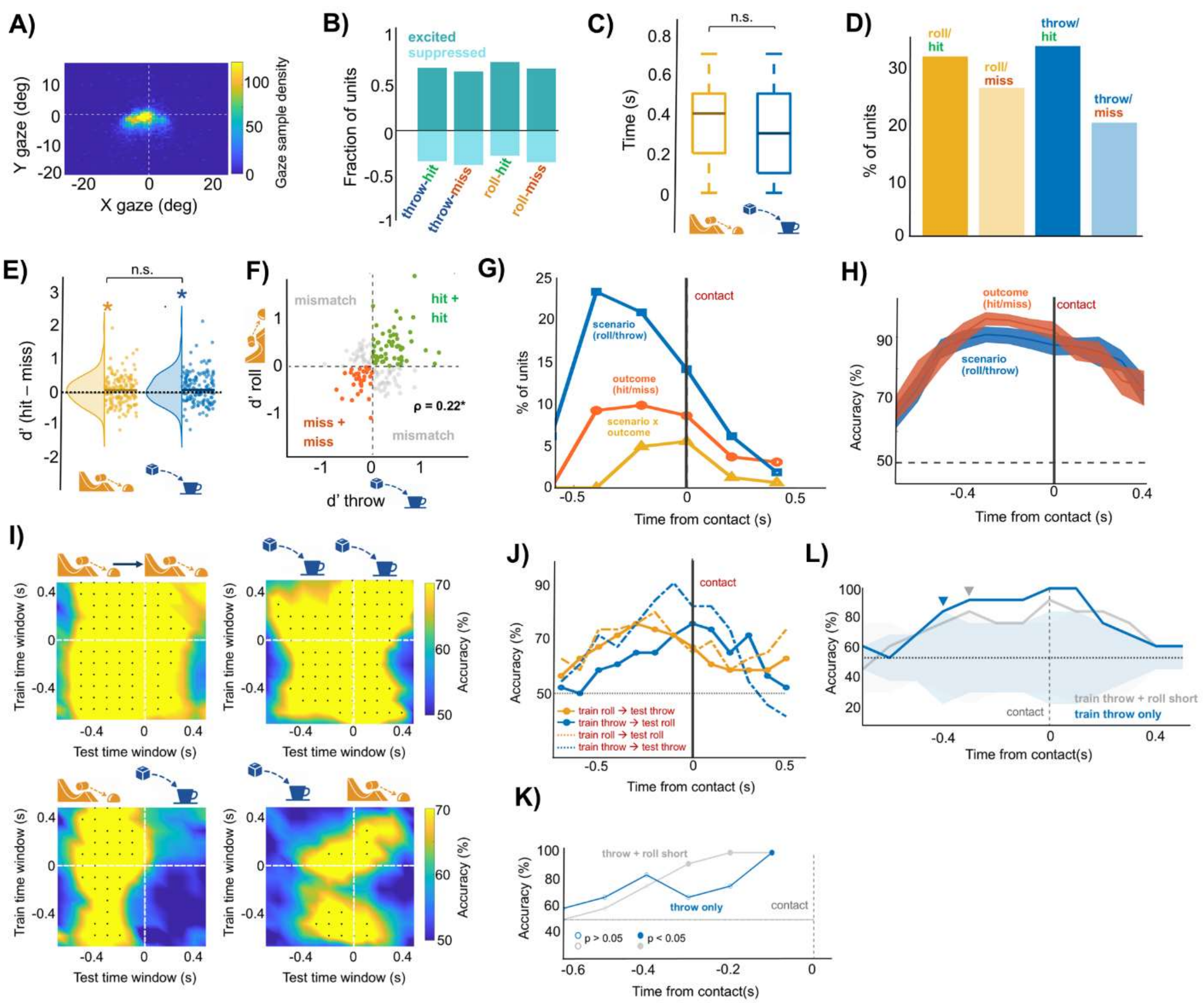


**(A)** Gaze distribution during the contact-prediction task. **(B)** Fraction of all recorded units showing excitation or suppression relative to baseline for each scenario and outcome. **(C to L)** Analyses from Figs. 3 and 4 were repeated using only well-isolated single units (SUA). **(C)** Response latencies for roll and throw trials (median, 400 and 300 ms before contact respectively). **(D)** Percentage of significantly responsive units for each scenario and outcome (roll-hit, 32.1%; roll-miss, 26.5%; throw-hit, 33.9%; throw-miss, 20.4%). **(E)** Distributions of hit-versus-miss d′ for roll and throw trials (median d′, 0.08 and 0.09, respectively). **(F)** Relationship between unit-level outcome selectivity across scenarios. **(G)** Time-resolved percentage of units showing significant main effects of scenario or outcome, or their interaction, in a two-way ANOVA. Peak proportions were 23.4% for scenario at −400 ms, 9.9% for outcome at −200 ms, and 5.5% for the interaction at 0 ms relative to contact. **(H)** Time-resolved decoding of scenario and outcome, reaching peak accuracies of 90.1% and 95.8%, respectively. **(I)** Temporal generalization of outcome decoding within and across scenarios. **(J)** Diagonal within- and cross-scenario outcome decoding extracted from (I). Peak accuracies before contact were 77.1% at −400 ms for roll→roll, 87.5% at −200 ms for throw→throw, 75% at −300 ms for roll→throw, and 64.6% at −300 ms for throw→roll. **(K)** Latency scan used to identify the precontact interval for cross-latency generalization. Decoders were trained on either throw and short-latency roll trials (gray) or throw trials alone (blue) and tested on held-out long-latency roll trials across successive precontact time windows. Filled circles indicate significant above-chance decoding after Bonferroni correction across tested windows ($P < 0.05$); open circles indicate non-significant decoding. Both training schemes first reached significance 100 ms before contact, defining the 200-ms interval centered on this time point (−200 to 0 ms) used for the cross-latency generalization analysis. **(L)** Cross-latency generalization control testing whether precontact outcome decoding generalized across differences in contact latency. Peak precontact decoding reached 83.3% at −300 ms for classifiers trained on throw + roll-short trials and 91.7% at −300 ms for classifiers trained on throw trials alone. Statistical procedures and plotting conventions are as in Figs. 3 and 4.

**Fig. S7. Physics selectivity in the contact-prediction session subset and outcome decoding in physics-preferring and control-preferring populations.**

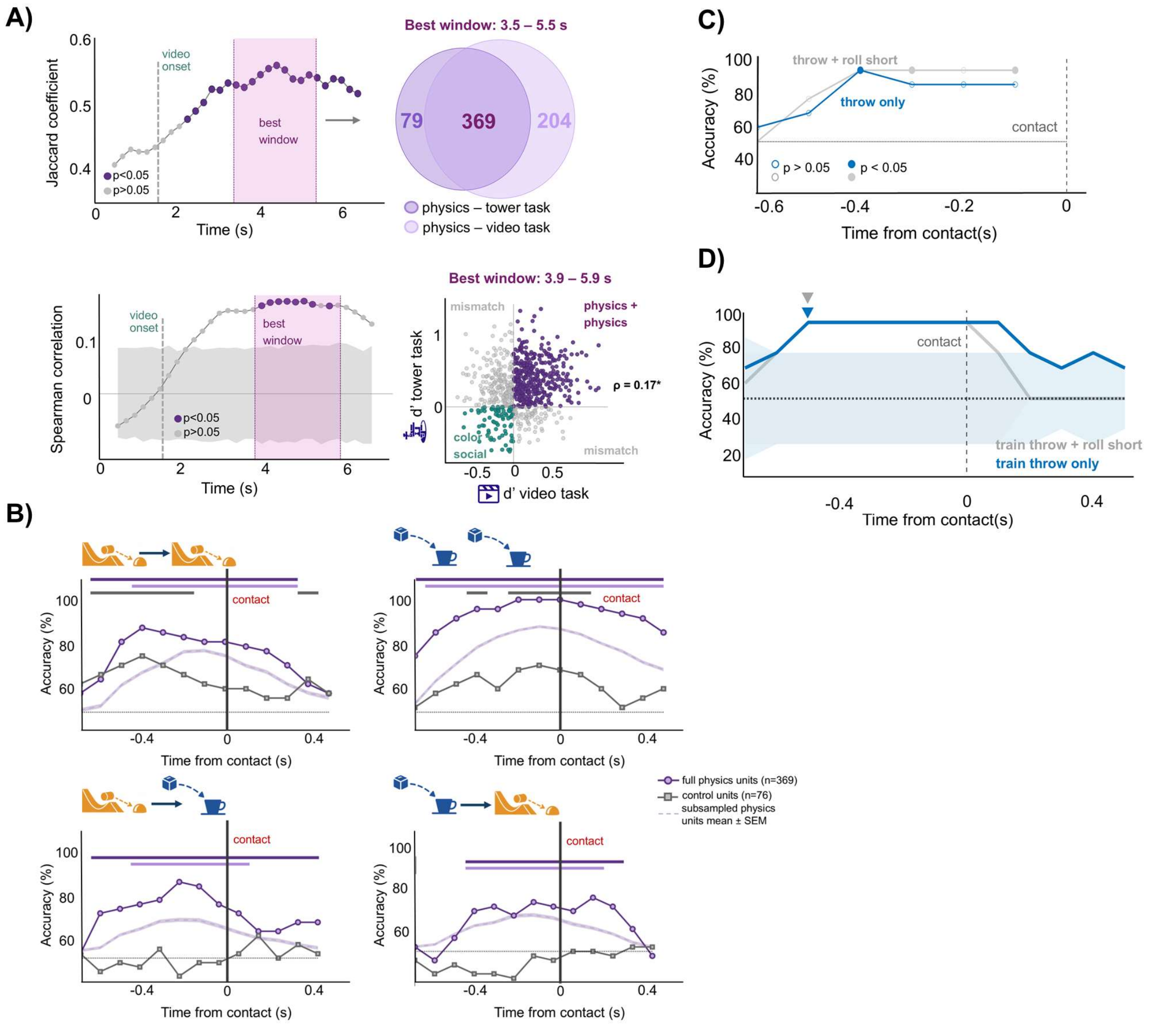


**(A)** Analyses from Fig. 2 repeated using the subset of recording sessions included in the contact-prediction experiment. Cross-task physics representations were assessed by the overlap of physics-selective units across the tower and video tasks (top) and correlation of unit-level physics selectivity across tasks (bottom). At the best-overlap window (3.5–5.5 s), 369 units were physics-selective in both tasks, with 79 selective only in the tower task and 204 only in the video task. Physics selectivity was significantly correlated across tasks within the best window (3.9–5.9 s;

Spearman ρ = 0.17, $P < 0.05$). Statistical procedures and plotting conventions are as in Fig. 2. **(B)** Within- and cross-scenario hit-versus-miss decoding using the full physics-preferring population (dark purple; n = 369), control-preferring population (gray; n = 76), and size-matched physics-preferring population (light purple; mean ± SEM across repeated subsamples). Diagonal decoding accuracies (matched train and test times) are shown for roll→roll (top left), throw→throw (top right), roll→throw (bottom left), and throw→roll (bottom right). Significant decoding emerged before contact in all populations within scenarios, but only in physics-preferring populations across scenarios. For roll→roll, decoding emerged at −600 ms in both the full physics-preferring (66.7% accuracy at onset) and control-preferring (64.6%) populations, and at −400 ms in the size-matched population (67.6%). For throw→throw, onset was −700 ms (75.0%), −600 ms (63.8%), and −400 ms (66.7%), respectively. For roll→throw, onset was −600 ms (70.8%) in the full physics-preferring population and −400 ms (63.5%) in the size-matched population; for throw→roll, onset was −400 ms in both populations (68.7% and 61.9%, respectively). Size-matched physics-preferring units showed significantly greater accuracy than control-preferring units at −300 ms for throw→throw, −400 ms for throw→roll, and −200 ms for roll→throw (one-sided normal-approximation tests, Bonferroni-corrected), with no significant differences for roll→roll. Horizontal bars indicate significant above-chance decoding; vertical lines mark contact (t = 0). Chance performance is 50%. **(C)** Latency scan used to identify the precontact interval for cross-latency generalization. To distinguish decoding aligned to the upcoming collision from activity related to elapsed time since video onset, decoders were trained on either throw and short-latency roll trials (gray) or throw trials alone (blue) and tested on held-out long-latency roll trials across successive precontact windows. Filled circles indicate significant above-chance decoding after Bonferroni correction across tested windows ($P < 0.05$). Both training schemes first reached significance 400 ms before contact, defining the 200-ms interval (−500 to −300 ms) used for the cross-latency decoding analyses in Fig. 4E. **(D)** Cross-latency generalization of outcome decoding. Decoders trained on either throw and short-latency roll trials (gray) or throw trials alone (blue) within the precontact interval identified in (C) were tested across the contact-aligned time course of held-out long-latency roll trials. Both decoders achieved 91.7% accuracy, demonstrating generalization across contact latencies and, for the throw-only decoder, across both scenario and latency. Triangles mark the first test time at which observed accuracy exceeded the pointwise permutation reference threshold; shaded regions show pointwise permutation-derived null intervals.

**Fig. S8. Action-intention responses and physics selectivity in the action-intention session subset.**

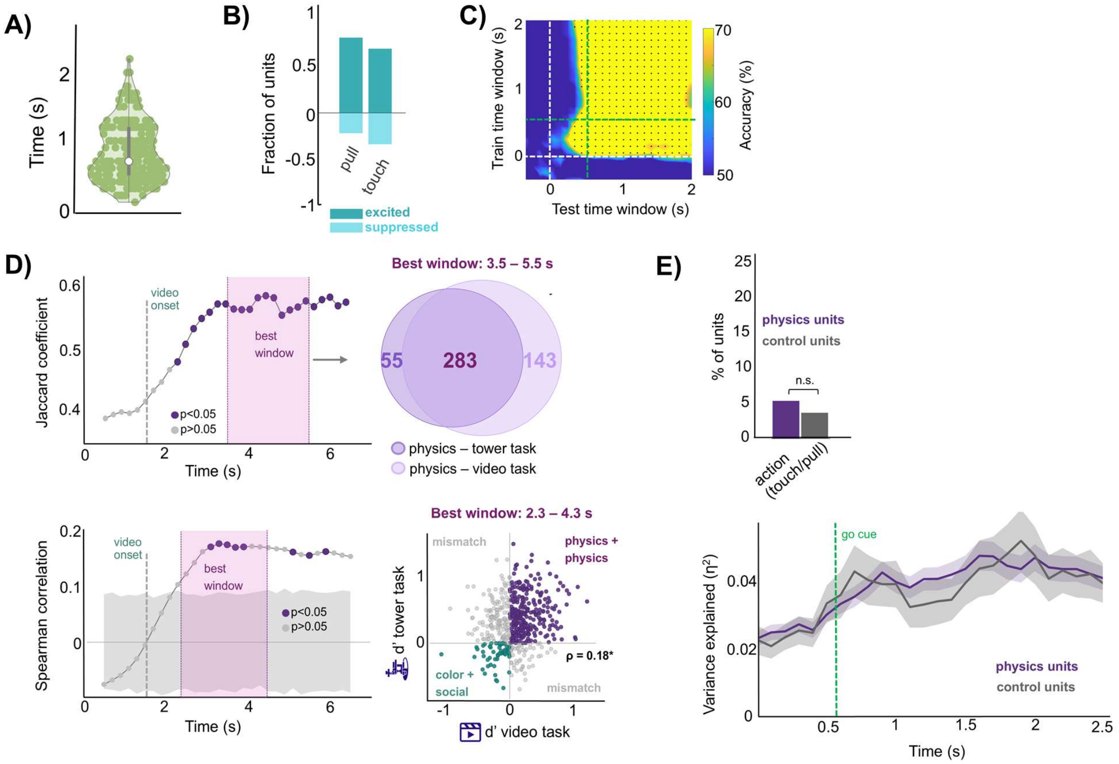


**(A)** Distribution of response latencies across the full population, defined as the time of first significant modulation relative to baseline. Points represent individual units. **(B**) Fraction of all recorded units showing excitation or suppression relative to baseline for pull and touch trials in the action-intention task. **(C)** Temporal generalization of touch-versus-pull decoding, with classifiers trained at each time window (y-axis) and tested across all time windows (x-axis). Color indicates decoding accuracy; black dots indicate significant train–test pairs. Dashed lines mark task events. **(D**) Analyses from Fig. 2 repeated using the subset of recording sessions included in the action-intention experiment. Cross-task physics representations were assessed by the overlap of physics-selective units across the tower and video tasks (top) and correlation of unit-level physics selectivity across tasks (bottom). At the best-overlap window (3.5–5.5 s), 283 units were physics-selective in both tasks, with 55 selective only in the tower task and 143 only in the video task. Physics selectivity was significantly correlated across tasks within the best window (2.3–4.3 s; Spearman $\rho = 0.18$, $P < 0.05$). Statistical procedures and plotting conventions are as in Fig. 2. **(E)** Percentage of physics-preferring and control-preferring units showing significant intended-action tuning (left), and variance explained ($\eta^2$) by intended action across the trial (right). Shaded regions indicate SEM. The green dashed line indicates the go cue. n.s., not significant.

**Fig. S9. Action intention responses and spatial distribution of physics- and action-related units.**

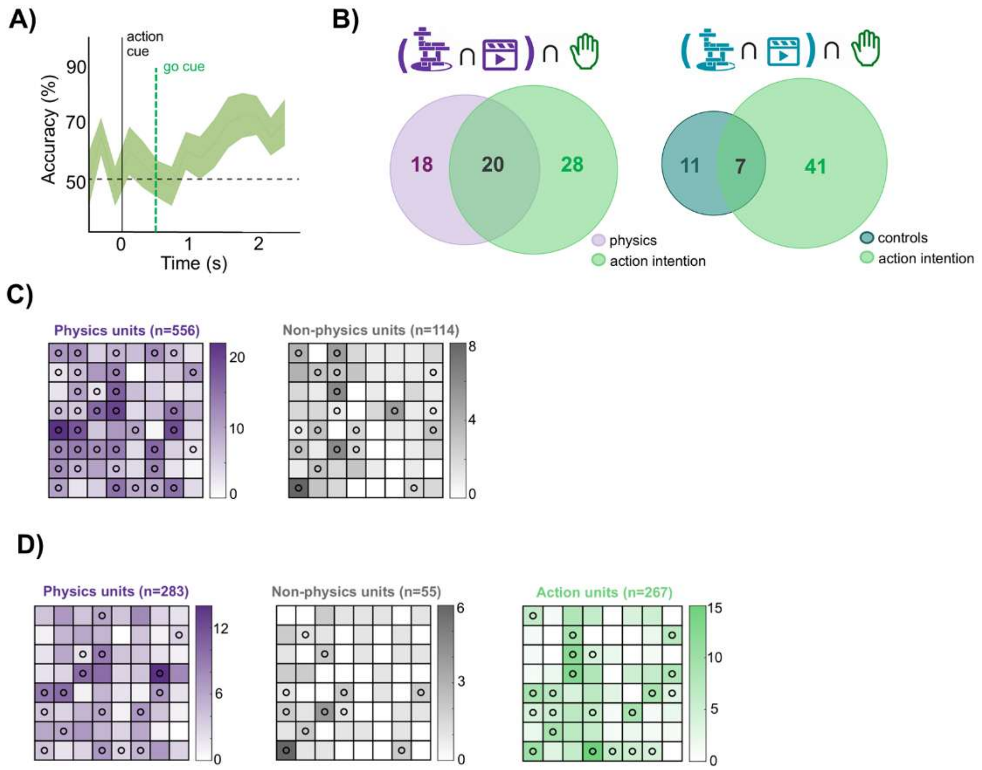


**(A)** Time-resolved SUA decoding of touch versus pull, aligned to the action cue, reaching a peak accuracy of 72.5%. The dashed green line marks the go cue, and the horizontal dashed line indicates chance performance (50%). **(B)** Overlap between physics-preferring and action-intention-responsive SUA (left) and between control-preferring and action-intention-responsive SUA (right). **(C)** Spatial distribution of physics-preferring (left) and control-preferring (right) units across the SPL electrode array using the full physics dataset. **(D)** Spatial distribution of physics-selective (left), non-physics (center), and action-intention-responsive (right) units across the SPL array for the subset of sessions containing the action-intention experiment. Color indicates the number of units recorded at each electrode; circles indicate electrodes containing at least one well isolated single unit from the corresponding population.

**Table S1. Recording sessions and experimental datasets in participant RD.**

| Session date | Tower | Video | Contact prediction | Action intention | SPL Units |
|---|---|---|---|---|---|
| 09/25/25 | ✓ | ✓ | ✓ | — | 81 |
| 10/23/25 | ✓ | ✓ | ✓ | — | 90 |
| 11/11/25 | ✓ | ✓ | ✓ | — | 99 |
| 11/25/25 | ✓ | ✓ | — | — | 99 |
| 12/02/25 | ✓ | ✓ | ✓ | — | 102 |
| 12/05/25 | ✓ | ✓ | ✓ | — | 95 |
| 05/05/26 | ✓ | ✓ | — | ✓ | 95 |
| 05/12/26 | ✓ | ✓ | — | ✓ | 89 |
| 05/19/26 | ✓ | ✓ | ✓ | ✓ | 84 |
| 05/21/26 | ✓ | ✓ | ✓ | ✓ | 86 |
| 05/26/26 | ✓ | ✓ | — | ✓ | 90 |
| 05/28/26 | ✓ | ✓ | ✓ | ✓ | 92 |

**Table S2. Recording sessions and experimental datasets in participant JJ.**

| Session date | Tower | Video |
|---|---|---|
| 10/03/25 | ✓ | ✓ |
| 10/22/25 | ✓ | ✓ |
| 10/23/25 | ✓ | ✓ |
| 10/28/25 | ✓ | ✓ |
| 11/06/25 | ✓ | ✓ |
| 12/09/25 | ✓ | ✓ |

**Table S3. Summary of recordings across experimental paradigms in participant RD.**

| Dataset | Sessions | All units | SUA |
|---|---|---|---|
| Tower / Video | 12 | 1,102 | 234 |
| Contact prediction | 8 | 729 | 162 |
| Action intention | 6 | 536 | 88 |

**Table S4. Stimulus-specific contact times used for contact aligned analyses.**

| Scenario | Setting | Contact time (s) | Stimuli |
|---|---|---|---|
| Roll | Indoor 1 | 0.90 | Hit 2; Miss 1 |
| | | 1.10 | Hit 1; Miss 2 |
| Roll | Indoor 2 | 0.82 | Hit 1–2; Miss 1–2 |
| Roll | Indoor 3 | 0.90 | Hit 2; Miss 1 |
| | | 1.45 | Hit 1; Miss 2 |
| Roll | Outdoor 1 | 0.82 | Hit 1–2; Miss 1–2 |
| Roll | Outdoor 2 | 0.89 | Hit 2; Miss 1 |
| | | 1.12 | Hit 1; Miss 2 |
| Roll | Outdoor 3 | 0.82 | Hit 1–2; Miss 1–2 |
| Throw | All settings | 0.80 | All hit and miss stimuli |
